# Understanding the dynamics of complex *in-situ* enzyme-catalyzed reactions undergoing mechanical stress

**DOI:** 10.1101/2023.10.30.564667

**Authors:** Baishakhi Tikader, Sandip Kar

## Abstract

Understanding the diversity in the enzymatic mechanism have utmost importance as it can temporally control the catalytic efficiency. Recent literature suggests that by influencing mechanical properties of hybrid materials, the catalytic efficiency of the enzymatic reactions can be altered significantly. Here, taking a computational and experimental approach, we have dig out the fate of the kinetics of enzyme reaction systems exhibiting relatively complex mechanism than usual Michaelis Menten kinetics involving multiple substrate/enzyme/enzyme-substrate complex. We have developed a numerical recipe improvising stochastic reaction-diffusion approach to explore the role of mechanical stress like compression-decompression cycle (C-D) on modulating the output of enzymatic reaction. The proposed methodology can be used as a theoretical tool to understand how to enhance the catalytic activity and setup appropriate reaction conditions by efficiently using the catalytic cycles. Hence, our methodology will be crucial to identify the most effective strategy to efficiently convert substrate into product in this type of mechanoresponsive materials, which will enable future development of cost-effective biomaterials. In future, the insights gained from this work may find enormous application in drug delivery, tissue engineering, bio-sensing and bio-catalysis.

## Introduction

Enzymes are natural biophysical catalysts, which crucially contributes to maintain different biological processes to sustain the continuity of the function of life. ^1,2^ Various biochemical processes starting from cellular membrane to tissue are known to regulate by enzymatic processes.^3^ The exact efficiency of these catalytic reactions depends on different external factors and can be fine-tuned by controlling various conditions such as pH, temperature, the amount of substrate and enzyme.^1,2^ The Classical Michaelis-Menten (MM) enzyme kinetics approximation has been extensively used to explore different aspect of the ensemble averaged enzymatic kinetic scenarios.^4–7^ However, at molecular level, studying the dynamics of enzymatic mechanism becomes more challenging as the spontaneous intrinsic fluctuations impart inherent stochasticity which influence the precision of the enzymatic processes.^7,8^ In this account, numerous theoretical and experimental techniques have been undertaken to investigate the distribution and conformational fluctuations associated with the enzymatic reactions.^4,6,7,9–12^

Recently, a new method has been developed to explore these biochemical reactions by integrating the functional enzymes in the form of nanoparticles as a mechanoresponsive biomaterials, and thereafter performing biochemical reactions in an efficient manner with these nano-enzymes.^13,14^ By creating such three-dimensional architecture, one can effectively regulate their physiochemical properties to control the catalytic output of such biochemical reactions. Thus, synthesizing and designing stimuli-responsive hybrid biopolymers, which can perform multi-compartment bio-catalyzed reactions has ignited a lot of interest in the field of enzyme catalysis.^13,15–18^ The synthesized macroporous hybrid biomaterials are extremely efficient for 2D cell culture and tissue development, hence can be introduced in tissue/organs for treating various diseases.^14,19–21^ Some of these nano-particle-based materials exhibit conducting properties and they have been used extensively as supporting electrical devices for different organs of the body (heart, brain, and skin).^19–28^ Complex enzyme-based networks that may exhibit competitive substrate binding or multiple intermediates, and cascade reactions are often found in natural biological processes.^29–31^ Understanding the complex in-situ catalysis reaction within such nano-materials can be an advanced step to optimally design such materials.^32–34^ On this account, investigating the inherent mechanism of catalysis within such materials and interpreting their kinetic progression can be a very crucial step to design and synthesize novel catalysts.

Consequently, understanding the catalytic property of these mechanoresponsive biopolymers is not straightforward. Stochastic intrinsic fluctuations influence the temporal turnover rate of these reactions.^35–39^ Along with this, due to the complex structure, most of these materials manifest compartmentalized architecture that brings spatial inhomogeneity into the system. Additionally, complexity arises, when the mechanism of the enzymatic processes involves multiple interconvertible conformational states, multiple substrates, multiple enzymes, or different binding strategies of substrates with the enzymes, which significantly complicates the in-depth understanding of the underlying kinetics.^38,39^ In this regard, predicting about the fate of the catalytic efficiency by performing mechanical stress on such biomaterial becomes extremely challenging. Experimental approach may not be sufficient here to unravel all aspects associated with the kinetic evolution of such catalysis reactions. Computational method can provide insights to dissect the inherent kinetics progression of compartmentalized biomaterial in the presence of external stimuli.

Herein, we intended to introduce a modified computational methodology (Stochastic reaction diffusion approach, SRDA) to capture the inherent inhomogeneity and stochasticity of any biochemical reaction (Module 1 and Module S1). Essentially, we have developed a new numerical recipe (Numerical method to incorporate C-D cycle, NMCD) to dissect the nature of the catalytic activity of such system under mechanical stress (such as compression-decompression [C-D] processes) (Module 2 and Module S2). The newly developed computational method (Method and Materials, Text S1) has been verified by investigating a known experimentally observed system and extended to different probable mechanistic mechanisms (Fig. S2). Our proposed numerical method enables to identify the regulating pathway, if the enzyme turnover rate or the fluctuating rate of intermediate states gets tuned by the [C-D] experiments. We speculate that this sort of investigation is critical to identify the most effective strategy to efficiently convert substrate into product, thus developing a new class of biocatalytic material in a cost effective manner. Our proposed methodology can be used as a theoretical tool to understand how to enhance the catalytic activity and setup appropriate reaction conditions by efficiently using the catalytic cycles. In future, the insights gained from this work may find enormous application in drug delivery, tissue engineering, bio-sensing and bio-catalysis.

## Methods and Material

### a. Efficient stochastic simulation algorithm of the reaction diffusion system (SRDA)

A stochastic description is needed to explain the spatiotemporal fluctuation in the concentration of chemically reacting species.^40–44^ Generally, the fluctuating dynamical evolution of different entities during the enzymatic reactions for a spatially homogeneous system can be easily formulated using Gillespie stochastic simulation algorithm (SSA).^45^ However, this assumption gets violated where the number of molecule is very low and the system exhibit a spatio-temporal inhomogeneity along with the stochasticity which is often observed inside the living cells. Additionally, the spatial inhomogeneity may also arise for the porous structure of a biomaterial. One of the ways to simulate this type of system is to partition the space into small homogenous spatial domains by assuming that the stochastic events can only occur within the same compartment and molecule can diffuse to its neighbouring compartments.^46^

Let us consider, *X*_j_ represents the number of molecule of a chemical entity in the j-th compartment. The entire computational domain in 1-dimension [0,L] has been divided into K compartments (the length of the compartment, *l* = *L*/*K*) as,

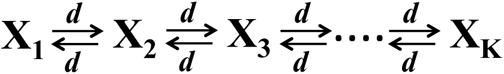

Where, 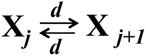 denotes that (i) 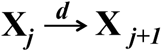 (Forward diffusion)

And (ii) 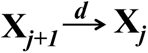 (Backward diffusion), *j* = 1,2,3,…, *K* [*d* = *D*/*l*^2^].

The diffusive property (denoted by diffusion constant, D) of the X molecule will depend on the temperature, viscosity of the solution and size of the molecule.

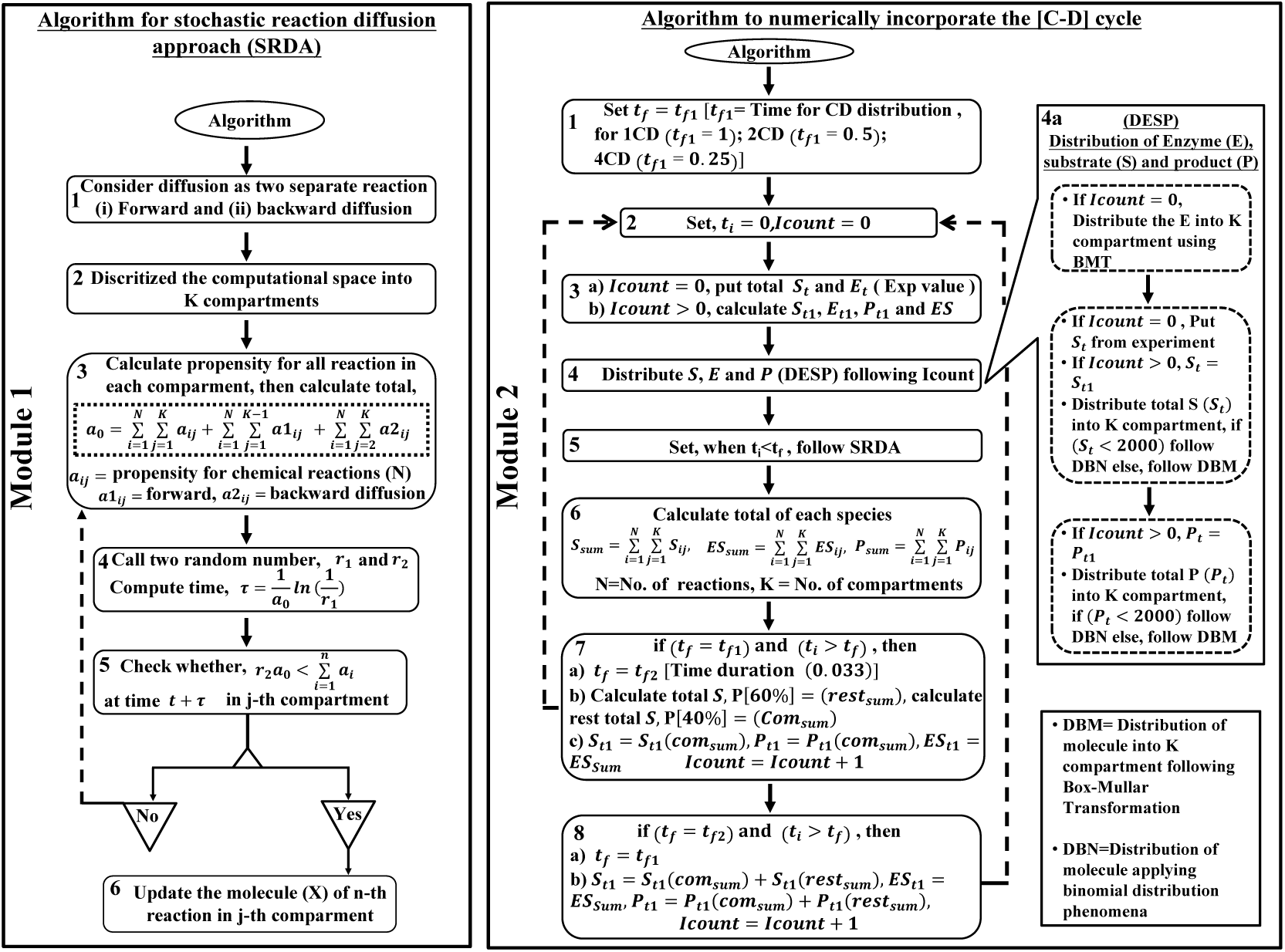

**Module 1.** Preliminary steps to simulate the stochastic reaction diffusion process (SRDA) for the compartmentalized system using a single loop. **Module 2.** Principle steps incorporated in the numerical method to mimic the effect of CD cycle on the rate of reaction of a biochemical reaction. The core module of the algorithm follows from step-1 to step-8 based on condition and coupled with two sub-modules (4a, 5a and 8a) necessary to complement the principle methodology. (The detail methodology has been incorporated in Text S1).

Erban *et al.* provided a methodology to compute the time evolution of X molecule by considering diffusion as a separate chemical reaction using the conventional SSA.^46^ However, their (SSA) based computational method formulates the update of the species in such a way that the calculation of propensity step repetitively appear after stimulating each elementary process thereby lengthening the existing numerical code. We have modified their approach to update all the reaction and diffusion events within a closed loop. A brief overview of the modified compartmentalized approach has been listed here.

1. Divide the diffusion as two different chemical reactions (forward diffusion and backward diffusion, Step-(1-2) of Module 1).
2. Calculated the corresponding propensity functions separately (Step-(3) of Module 1).
3. Afterwards, generate two random numbers for finding the probability of occurring n-th reactions in between time *t* and (*t* + *τ*) (Step-(4-5) of Module 1).
4. Update the number of molecule for n-th reaction in j-th compartment (Step-6 of Module 1).
5. Continue with step-1 upto desire time.

Thus, our proposed methodology (See Text S1 in supplementary for detail) helps to simulate the stochastic reaction diffusion process with better efficiency and reduces the computational cost.

### b. Methodology to incorporate the compression-decompression phenomena in existing SRDA algorithm (NMCD)

1. In this work, we have proposed a generalized numerical recipe (Module 2 and Text S1) to validate the increase in the product formation when an enzymatic reaction of a spongy biomaterial has been performed under mechanical stress. We have developed the new numerical methodology (Module 2) to mimic the compression-decompression [C-D] experiment performed by mehek et *al.*^13^ A brief overview have been provided to summarize the proposed methodology. Starting with initial conditions, we perform six steps at time t.
2. Consider two separate time scale, define the final time (*t*_*f*_) as either (*i*) *t*_*f*1_ (the time span for the normal chemical reactions) or (*ii*) *t*_*f*2_ (the time duration required for performing the mechanical operations due to [C-D] cycle) (Step-1 of Module 2).
3. Set initial time, *t*_i_ = 0 (with a counter *Icount*(= 0)) (Step-2 of Module 2).
4. Redistributed the molecule either using the Box-Mullar transformation or binomial distribution, after each [C-D] cycle (Step-(4-4a) of Module 2).
5. Simulate the system using SRDA protocol for *t*_i_ < *t*_*f*_ (Step-5 of Module 2).
6. Keep tracks of the total of every species and calculate the total amount of the substrate and product in similar proportions separately and update the *Icount* (Step (6-8) of Module 2).
7. Continue with step-1 upto the desire time (in our case at 25 minutes) (Step-8a of Module 2. Additionally, we constructed a separate module and coupled with the present methodology to incorporate the alteration in the reaction rate due to the [C-D] cycle (See Text S1 in supplementary for detail).

In summary, the primary goal of this work is to explore different probable enzymatic reaction schemes occurring within these spongy materials under the influence of mechanical stress. Our strategy is to modify the existing SRDA to make it an efficient algorithm, such that the new methodology incorporates the [C-D] cycle phenomena (NMCD) along with the SRDA. Fig 1 represents generalized schematics to capture this kind of enhancement in the rate of formation of the product during [C-D] cycle using our NMCD method.

**Fig. 1.**
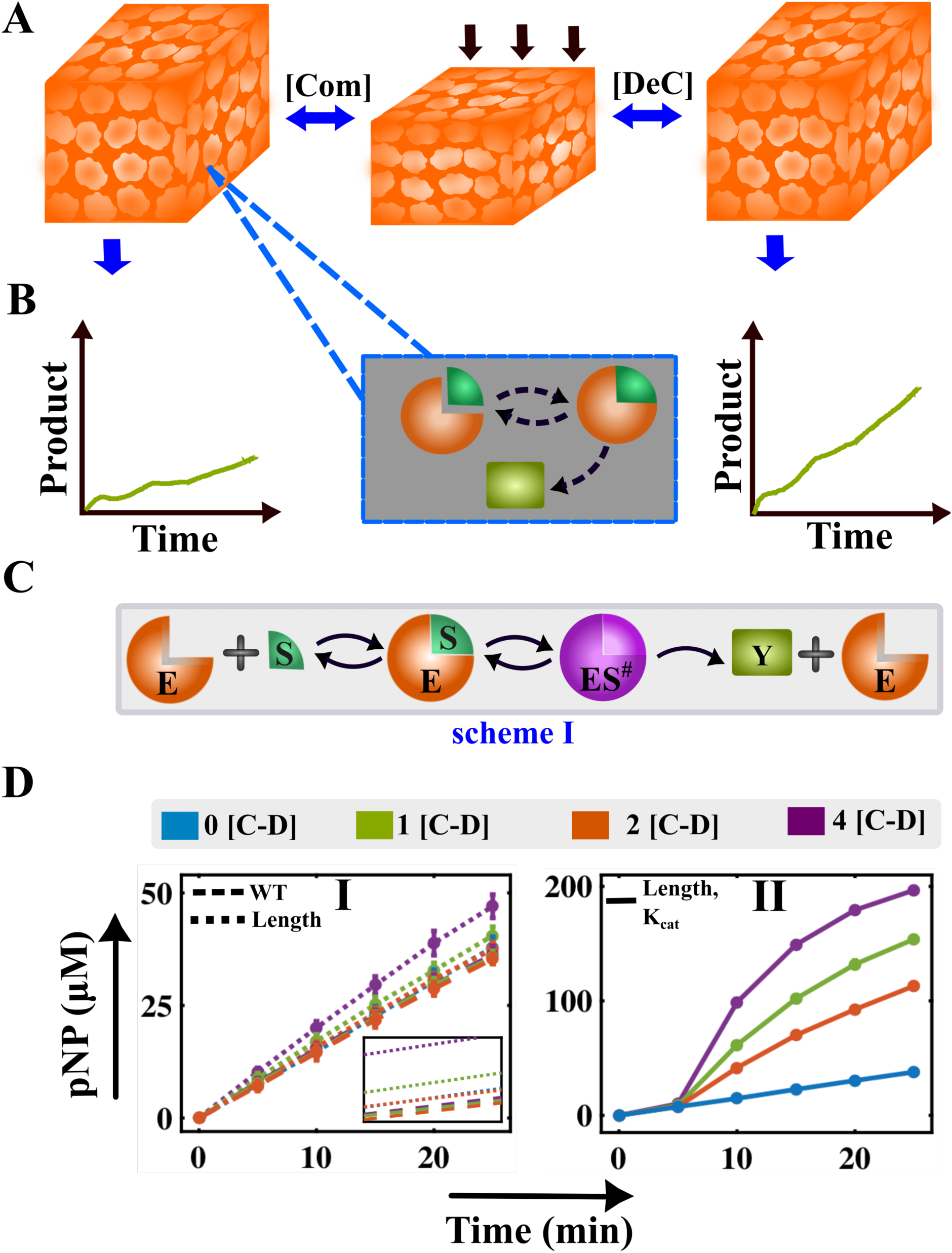
Understanding the product formation kinetics of biochemical reactions following scheme-I within a biomimetic scaffold undergoing [C-D] cycles. **A)** Schematic of an enzyme-catalyzed biochemical reaction occurring inside a compartmentalized spongy biomaterial undergoing compression [**C**] – decompression [**D**] cycles. **B)** Probable product formation profiles of enzyme-catalyzed reactions which can involve multiple enzymes or substrates or intermediate conformational states or products. **C)** Proposed mechanism of scheme-I, having two interconvertible non-equivalent conformational states (ES and ES^#^). **D) I)** Enhancement of the product formation (dashed lines) for the increment in the binding rate of the enzyme with the substrate (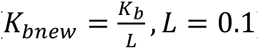) due to the compression of the length (Text S3D), while performing [C-D] cycles compared to the CT case, (dotted lines, when the rate constant has not been altered, Table S2). **II)** Validation of the experimental observation of the enhancement of the product formation upon assuming that [C-D] cycles (0, 1, 2, and 4) fine-tune the catalytic efficiency of the enzyme (Text S(3-4)) for enzyme-catalyzed reaction considered in the schematic-I. In each case **D(I-II)**, the solid lines represent the average trajectory of 5 runs with the error bars.

Thereafter, we successfully implemented this technique for different reaction schemes (Fig. S2, Scheme-(I-V)) to analyze the outcome of the enzymatic reactions undergoing mechanical stress. In this project, we consider enzymatic reaction systems exhibiting relatively complex mechanism than usual MM kinetics by either having two interconvertible conformations leading to one product (Scheme-I), competitive product formation from two different intermediate complex (Scheme-(II-III)), an enzyme targeted by two substrates forming two consecutive products (Scheme-IV), and a substrate encountering two enzymes to produce two separate products (Scheme-V).^47,48^ From a catalysis perspective, we believe that this proposed method can suggest a new strategy to rationally design such biomaterial that can enhance the reproduction of the complex biochemical system exhibiting such mechanoresponsive property.

## Result and discussions

### Dynamics of the product formation for reaction mechanism having multiple conformational states (scheme-I)

To establish any theoretical methodology, having a close correspondence with experimental observation is imperative. In order to do that, first, we have studied a very simple mechanism following scheme-I. Here, the substrate (*S*) reversibly binds with the enzyme (*E*) and sequentially produces two interconvertible intermediate states, *ES* and *ES*^#^(*ES*^#^is another conformational state of *ES*, where either *E* or *S* changes their conformational state), respectively. *ES*^#^irreversibly converts into the product (*Y*) and regenerates the enzyme (*E*) (Fig. 1C, S1A). Mehek *et al.* has synthesized a silica/silk nanoparticle-based enzyme-polymer surfactant porous scaffold, which can catalyze the hydrolysis of p-nitrophenyl phosphate (pNP).^49^ The enzymatic reaction happening within the scaffold has a resemblance with the proposed scheme-I. Hence, the proposed scheme-I has been applied (Fig. 1C, Text S1) to validate this experimental data of the hydrolysis reaction under the [C-D] cycles.^49^ Here, one extra layer of complexity in the mechanism has been added (considering two intermediate states) in comparison to the simple MM mechanism. The proposed kinetic scheme (Text S4, Table S(1-4)) has been simulated under the appropriate conditions using SRDA (Test S4) in absence of the compression-decompression cycles (0 [C-D]). It has been found that scheme-I is able to represent the rate of formation of the product (Fig. S2A) satisfactorily for the 0 [C-D] cycles as observed experimentally.

Mehek *et al.* have observed that the product formation rate increases with increasing frequency of [C-D] cycles (1, 2 & 4). The NMCD method has been implemented to mimic this experimentally observed higher rate of product formation. Surprisingly, the proposed methodology is unable to capture the enhancement of the rate of the product formation (Fig. 1DI, WT, dashed lines) under [C-D] cycles. It has already been observed in the previous chapter that for this kind of biomaterial, where the species get re-distributed more or less uniformly during each [C-D] cycle, the diffusion has a very insignificant role in controlling the output of the respective biochemical processes (Fig. S2B). Thus, the effect of diffusion on the reaction rate has not been explored for other schemes (II-VI) as well. Instead, an appropriate algorithm has been implemented into the existing methodology (Text S1(C-D)) to distribute enzyme (at the beginning of each simulation since the enzymes are embedded in the spongy biomaterial), substrate, and product (at the end of each decompression) to mimic the experimental scenario as observed for these synthesized biomimetic materials.

Literature survey suggests that the bimolecular reaction rate rises up as the length of these biomaterials gets decreased due to compression. Incorporation of this fact (*L* (*new*) = *L* /10 for all [C-D] cycles) into the existing method results in a steady increase in the production rate (Text S2D), Fig. 1DI, length, dotted lines) with 1, 2, and 4 [C-D] cycles. However, it is unable to capture the sharp rise as observed in the experiment.^49^ A robust increase in the rate of the formation of product (*Y*) with ascending [C-D] cycles pinpoint a significant alteration of other reaction rates occurring during the [C-D] activity. Thus, the probable variation of the various reaction rates during the numerical implementation of [C-D] events has been explored. It has been found that the modification in catalytic rate (*k*_*x*_) or the interconversion rate (*k*_i1_) of the enzyme-substrate complexes produce the proper temporal increment in the product formation with alteration in the [C-D] cycles, which nicely corroborates with the experimental observations (Fig. 1DII). It can be concluded that the enhancement in the product formation for this particular material is arising mostly from the direct modulation of the internal fluxes (either catalytic rates or interconvertible rates) mediated by mechanical stress. This encourages me to explore the fate of the kinetics of the product formation of the next complex enzymatic reaction (Fig. S1B) occurring inside the spongy biomimetic material. Here, we have proceeded by sequentially adding the complexity to the current reaction mechanism (scheme-I) to understand how [C-D] cycles influence the complex enzymatic reaction pathway in a similar manner (as observed for scheme-I). The computational study may be useful in predicting the most effective way to obtain a specific product in a reasonable amount under the hypothesis that the [C-D] cycles may influence reaction propensities in a specific manner.

### Intermediate conformational states control the dynamics of the multiple product formation (scheme-II)

The next probable enzymatic scheme (scheme-II) that has been considered is quite similar to that of scheme-I. In this scheme (Fig. 2AI, Table S(4-6)), another product formation *X* has been assumed in such a way that the extent of the product *X* formed from the *ES* complex will compete with the formation of product *Y* from another conformer (*ES*^#^). Here, the primary goal is to monitor whether it is possible to increase the product formation towards any desired direction by employing the [C-D] cycles. It has been considered that the kinetic rate constants of current reaction scheme-II (Fig. S1B, Table S6) can lead to three different scenarios under three different parametric conditions; (i) the amount of the product *X* and *Y* formed are almost similar (*C*_1_), or (ii) the amount of product *X* can be moderately higher than the production of *Y* (*C*_2_), and (iii) the amount of product *Y* can be comparably higher than the product *X* (*C*_3_) (Fig. 2AII) under 0 [C-D] cycle.

A probable generalized free energy diagram has been depicted in Fig. 2AIII schematically to represent different scenarios (*C*_1_, *C*_2_ and *C*_3_) of the enzyme-catalyzed reaction.^50^ Here, searching specific microscopic reaction propensities are of keen interest, which can get fine-tuned due to the employment of the [C-D] cycles and can drive the evolution of *X* or *Y* under different conditions (*C*_1_, *C*_2_ and *C*_3_). As observed for scheme-II, only incorporation of the [C-D] cycles (1,2 & 4 [C-D]) during simulation is ineffective to alter the product formation (CT, dashed lines, Fig. S3). Though, an effective enhancement in the emergence of the product has been noticed (Length, dotted lines, Fig. S3), after introducing the increment in the bimolecular rate arising from the alteration of the length of the material (Text S3D, *L* (*new*) = *L* /10, *k*_*b*_ (*new*) = 10 × *k*_*b*_ during [C-D]) events. Investigation of scheme-I has revealed that the [C-D] cycles may influence the catalytic rates or the conversion rates of intermediates and alter the amount of product formation. In this scheme-II, the catalytic rate (*k*_*x*_) controls the production of *X*, hence, under any specific parametric condition, if the [C-D] cycles are able to enhance *k*_*x*_ that may result in the acceleration of the formation of *X*. However, the modulation of the catalytic rate (*k*_*y*_) through [C-D] cycles, fails to aid the production of *Y* (Fig. S4). At this point, hypothetically it has been considered that the [C-D] cycles may fine-tune the conversion rate from *ES* to *ES*^#^ (*k*_i1_), which can eventually alter the creation of *Y*.

In the case of *C*_1_, where both the products can be generated from scheme-II under parametric conditions (Table S6) in almost equal amounts (blue lines, Fig. 2AII, the amount of product *X* or *Y* can be altered easily by implementing [C-D] cycles via changing the respective reaction fluxes (*k*_*x*_ or *k*_i_). It has been predicted from the model simulations that if the *k*_*x*_ get escalated almost ∼100 (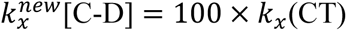) folds due to [C-D] activity, the propensities of the formation of *X* shoots up (Fig. 2BI) with simultaneously decreasing the yield of product *Y* (Fig. 2BII) compared to the control CT (Fig. S5A(I-II)). On the contrary, if the [C-D] cycle increases the intermediatory conversion rate (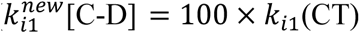), it helps to accelerate the formation of *Y* (Fig. 2BIV) and results in a moderate decrease in the production of *X* (Fig. 2BIII) compared to CT (Fig. S5A(III-IV)).

For the case *C*_2_, the parametric conditions (Table S6) lead to the formation of *X* more (red lines, Fig. 2AII) for the 0 [C-D] cycle in comparison to *Y*. Thus, the enhancement of *k*_*x*_ due to the [C-D] cycles (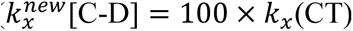) causes a monotonic rise in the amount of *X* (Fig. 2CI, S5BI) and creates a marginal reduction in the production of *Y* compared to CT (Fig. 2CII, SBII) as expected. In contrast, if the [C-D] cycle alters the conversion rate *k*_i1_, the product of *Y* increases in comparable amount (∼6 times, Fig. 2CIV, S5BIV) and reduces the formation of *X* to moderate extent compared to CT (Fig. 2CIII, S5BIII). Under *C*_3_ condition, simulating the scheme-II generates a significant amount of *Y* compared to *X* (ruby lines, Fig. 2AII) under 0 [C-D] situation. Even though the reaction propensity is significantly biased towards the creation of *Y* (for the 0 [C-D]), the increment in the catalytic fluxes (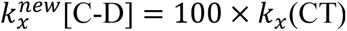) due to the [C-D] activity can amplify the production of *X* (Fig. 2DI) considerably by lowering the production of *Y* (Fig. 2DII) compare to CT (Fig. S5C(I-II). Due to the implementation of the [C-D] cycles, a faster rate of conversion (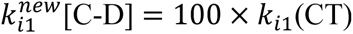) helps to accumulate *Y* in an appreciable amount and decreases the production of *X* (Fig. 2D(III-IV), S5C(II-IV)) as expected. A probable representative free energy diagram for the respective product formation under the alteration of these rates has been schematically depicted in Fig. S6.

**Fig. 2.**
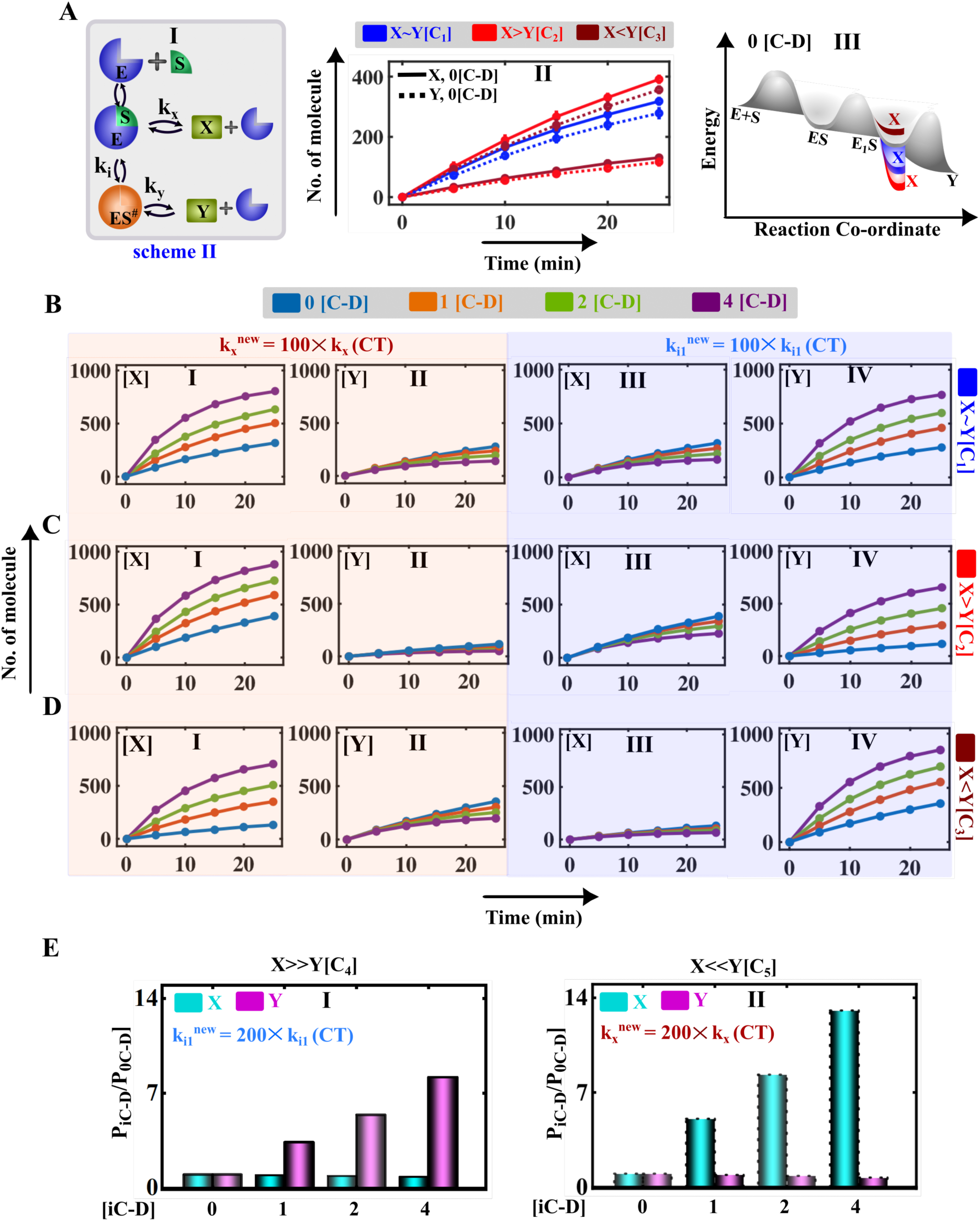
Kinetic study of the enzyme-catalyzed reaction producing two consecutive products (scheme-II) in the absence and presence of [C-D] cycles. **A) (I)** Scheme-II describes an enzymatic reaction that gives two products *X* and *Y*. Time course simulations of the **(II)** product*X* and **(III)** product *Y* under three different scenarios (*X*∼*Y*, *C*_1_), (*X* > *Y*, *C*_2_) and (*X* < *Y*, *C*_3_) for 0 [C-D] cycles (corresponding parameter set is provided in Table S1). **IV)** Probable free energy profile diagram for the catalytic reaction following scheme-II under *C*_1_, *C*_2_, and *C*_3_ conditions.^50^ The amount of product *X* (**BI** for case *C*_1_, **CI** for case *C*_2_, and **DI** for case *C*_3_) and product *Y* (**BII** for *C*_1_ case, **CII** for *C*_2_ case, and **DII** for *C*_3_ case), if the [C-D] cycles accelerate the catalytic rate *k*_*x*_100 times compared to CT. The amount of product *X* (**BIII** for case *C*_1_, **CIII** for case *C*_2_, and **DIII** for case *C*_3_) and product *Y* (**BIV** for *C*_1_ case, **CIV** for *C*_2_ case, and **DIV** for *C*_3_ case), when the [C-D] cycles fine-tune the conversion rate as (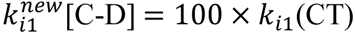). The product formation for scheme-II depicts two extreme scenarios, where *X* >> (solid lines, *C*_4_) and *X* << *Y*(dotted lines, *C*_5_), without any compression-decompression experiment (0 [C-D] cycle) using the parameter set provided in (Table S1). The relative increment of the level of products *X* **(EI)** and *Y* **(EII)** (where *P*_i*CD*_/*P*_0*C*D_, *i* = 1, 2, 4) by altering the *k*_i1_ and *k*_*x*_ (200 times) under different [C-D] cycles for the*C*_4_ and *C*_5_ cases, respectively.

This observation suggests that any biochemical reactions following the mechanism depicted in scheme-II can produce a higher level of *Y* under particular conditions. Even, in these conditions, it is possible to drag the dynamics towards the production of *X* in a considerable amount by applying mechanical stress. However, once the system is tilted towards the formation of *X*, the alteration in the reaction mechanism allows the accumulation of *Y* comparatively in a lesser amount. To make the observation more noticeable, scheme-II has been further analyzed under two extreme scenarios *C*_4_ and *C*_5_. For *C*_4_ case, though the system is significantly titled towards the production of *X* (Fig. S7A) under the 0 [C-D] condition, still the production rate of *Y* can be enhanced nearly 8-fold if the conversion rate (*k*_i1_) get altered appreciably due to the [C-D] cycles (Fig. 2EI, S7(C-D)). Whereas, for *C*_5_ case, the system is considerably biased to create more *Y* compared to *X* under the 0 [C-D] condition (Fig. S7B). Here, if the [C-D] cycles are able to alter *k*_*x*_ significantly, it is possible to produce more *X* compared to CT (almost 15 folds, Fig. 2EII, S7(E-F)).

This observation involving scheme-II signifies that the product formation ability can be altered significantly by employing mechanical stress in the form of the [C-D] cycles. These observations suggest that the specific design of the biomimetic material and the nature of the substrate allow the alteration of particular microscopic steps involved in the enzymatic reaction (by modifying some specific chemical interaction or some related effect) during [C-D] cycles. This can help to modify the extent of the output of the corresponding reaction system in any desired path by designing an appropriate experimental setup. This idea encourages me to understand the role of the [C-D] cycles in more complex enzymatic reactions than scheme-II, where the influence of the [C-D] cycles on the intermediate complexes can be studied extensively.

### Substrate transduced intermediate conformer significantly modulates the kinetics (scheme-III)

The analysis of scheme-II reveals that the interconvertible conformers play a pivotal role in rationalizing the enzyme turnover rate under [C-D] cycles. Hence, another level of complexity has been added by introducing another substrate (*S*_2_) in scheme-II. In this scheme (scheme-III) (Fig. 3AI, S1C, Table S(7-9)), an additional substrate (*S*_2_) binds with the enzyme-substrate complex *ES*^1^ directly to form a hetero-trimeric complex *ES*^##^that eventually gives rise to *Y*. The product *X* originates immediately from the enzyme-substrate complex *ES*^1^. Thus, the formation of product *X* competes with the creation of the *ES*^##^complex. As shown for the scheme-II, scheme-III can dynamically exist in three different states (*C*_1_, *C*_2_ and *C*_3_, Fig. 3A(II-III)) under three different parametric conditions (Table S9), and can lead to similar observations upon altering the length of the material as observed for scheme-II (Fig. S8). For all the cases, it has been assumed that the [C-D] cycles change the rates of the reaction (either catalytic or transition rate) as the length of the material is altered. When the reaction has an equal probability to form either product *X* and *Y* (blue lines, Fig. 3AIII, alteration of distinct microscopic rates (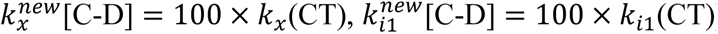) during the [C-D] cycle enhances the *X* or *Y* in proportionate amount (Fig. 4b.3B(I-IV), S9A(I-IV)) compared to CT as observed in case of scheme-II.

**Fig. 3.**
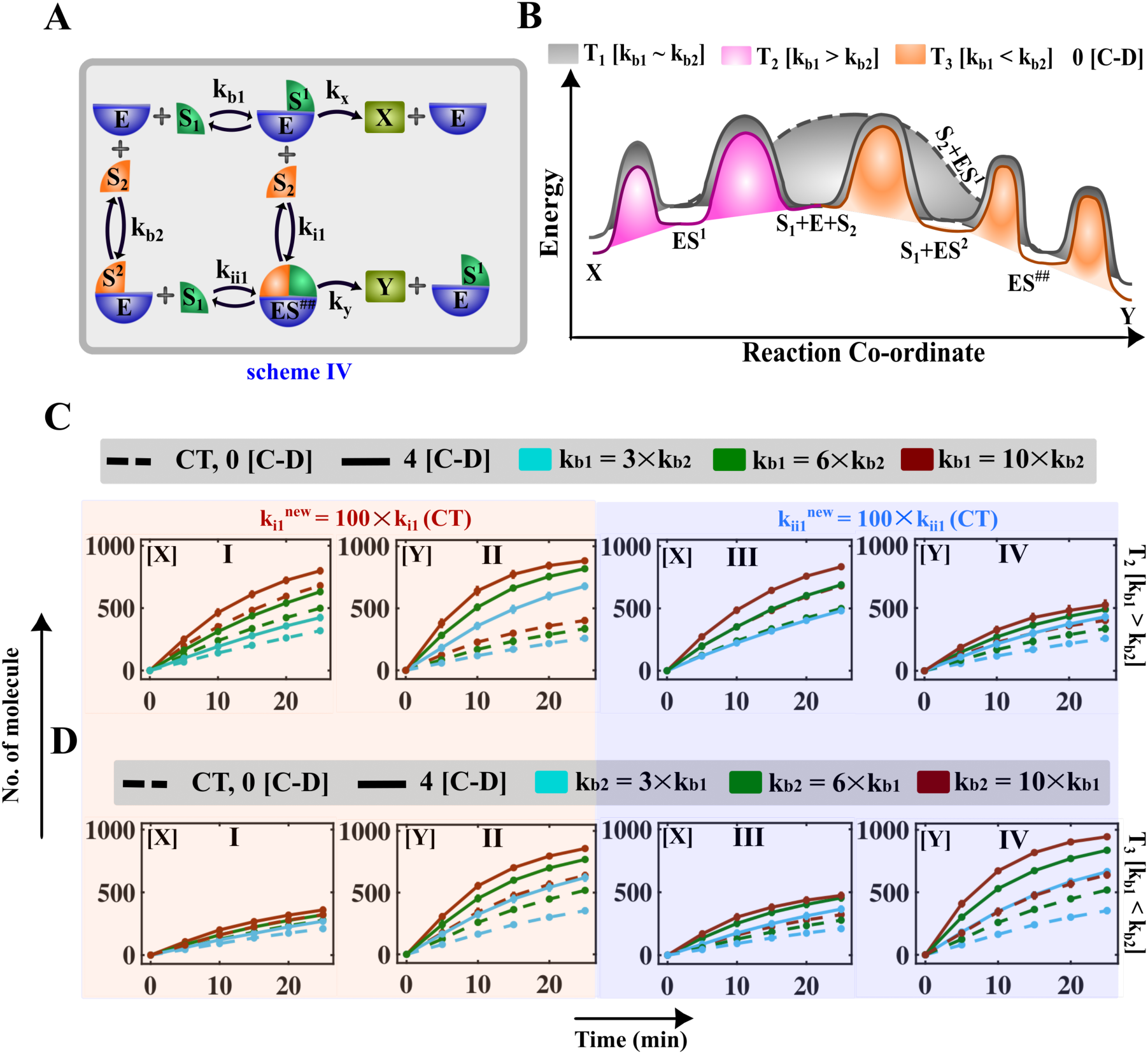
Fine-tuning of the formation of products **X** and **Y** involved in the biochemical reaction following scheme-IV is possible by employing C-D cycles. A) The probable mechanistic pathway of a biochemical reaction is depicted in scheme-IV. B) One of the probable ways to represent the free energy diagram of catalysis reactions undergoing three different conditions based the binding rate of two substrates has been fined-tuned such that (1) (**T**_**1**_) both product (**X**) and (**Y**) can reproduce either in a comparable amount or (2) (**T**_**2**_) the formation of product (**X**) is higher than the product (**Y**) or vice-versa (**T**_**3**_) for 0 [C-D] cycle. C) Simulated time profiles of I) product (**X**) and II) product (**Y**), when the conversion rate (**k**_**i1**_) has been enhanced 100 times and time profiles of III) product (**X**) and IV) product (**Y**), altering the conversion rate (**k**_**ii1**_) 100 times during [C-D] cycles (4 [C-D]) compared to CT in case **T**_**2**_. The simulation has been performed by increasing the binding rate as **k**_*ii***1**_ = **3** × **k**_**b2**_ (cyan color), **k**_**b1**_ = **6** × **k**_**b2**_ (green color) and **k**_**b1**_ = **10** × **k**_**b2**_ (maroon color) keeping all other rates constant same as in Table S9. D(I-IV)) A similar analysis has been performed taking **k**_**b2**_ = **3** × **k**_**b1**_ (cyan color), **k**_**b2**_ = **3** × **k**_**b1**_ (green color) and **k**_**b2**_ = **10** × **k**_**b1**_(maroon color). The dotted lines represent the simulated curves for CT (Table S11). The dashed lines represent the product formation for the increment of bimolecular rate (**k**_**b1**_/**L**, **k**_**b2**_/**L, L** = **0. 1**) due to the compression of length (**L**) while performing 4 [C-D] cycles.

**Fig. 4.**
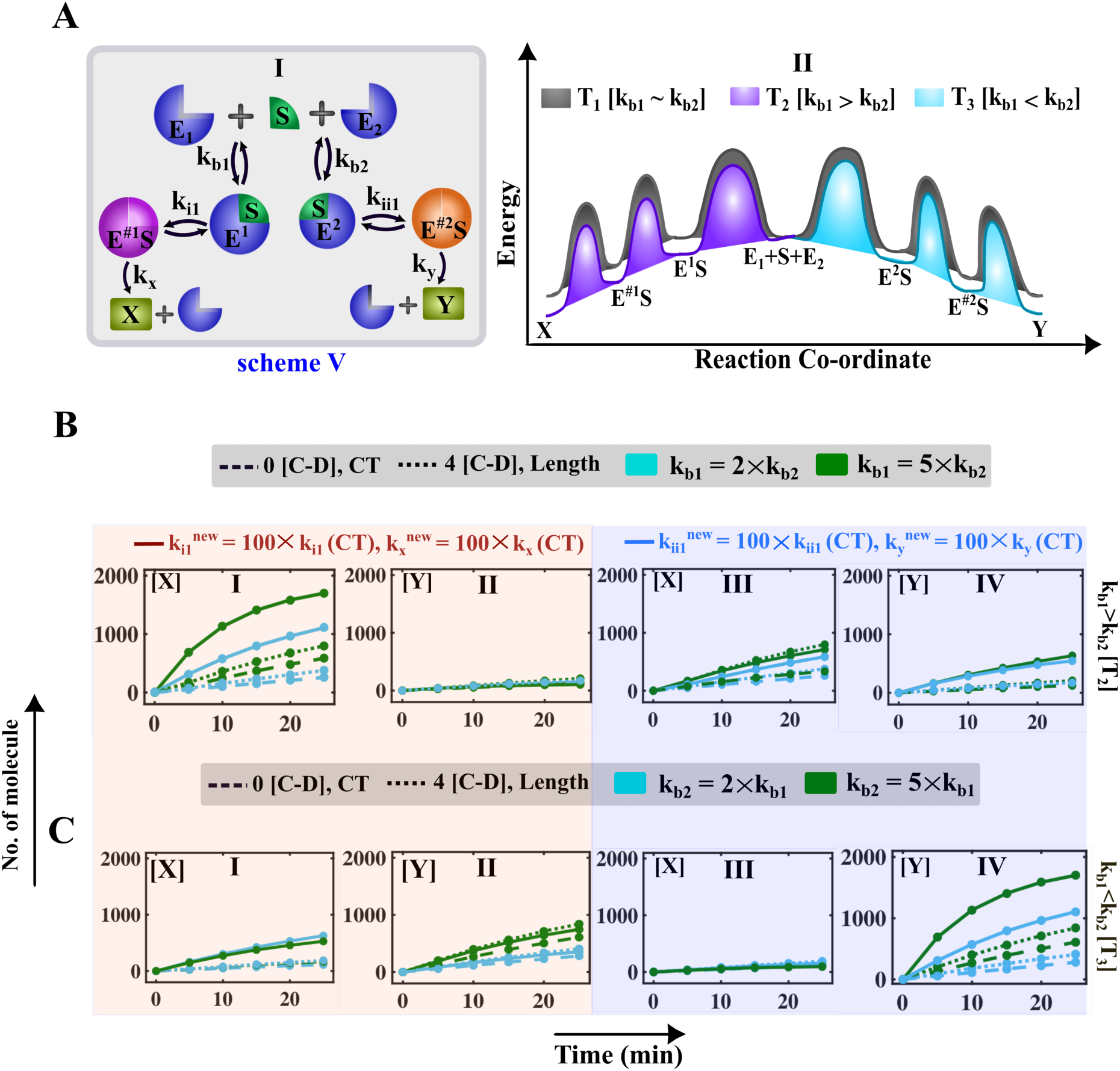
Fine-tuning of the formation of products *X* and *Y* produced from biochemical reaction following scheme-V due to [C-D] cycles. **AI)** The biochemical reaction scheme has been depicted in scheme V. The enzymes *E*_1_ and *E*_2_ consecutively bind with the same substrate *S* and produces two different enzyme-substrate complexes *E*^1^*S* and *E*^2^*S*, which can convert into another intermediatory enzyme-substrate complex *E*^#1^*S* and *E*^#2^*S*. The intermediate state *E*^#1^*S* can give product *X*, and the intermediate state *E*^#2^*S* can give product *Y* and regenerate the enzyme *E*_1_ and *E*_2_. **AII**) One of the probable energy diagrams of the proposed mechanism for the three conditions **T**_1_, **T**_2_, and **T**_3_. **B)** For case **T**_2_, the simulated time profiles of **BI)** product (*X*) and **BII)** product (*Y*), when [C-D] events (4 [C-D]) fine-tune the intermediate rate *k*_i1_ and the catalytic rate *k*_*x*_ collectively (solid lines). Time evolution of **BIII)** product (*X*) and **BIV)** product (*Y*), when [C-D] event (4 [C-D]) alters the intermediate rate *k*_ii1_ and the catalytic rate *k*_*y*_ collectively (solid lines) compared to CT. The simulation has been performed by increasing the binding rate as *k*_*b*1_ = 2 × *k*_*b*2_ (cyan color) and *k*_*b*1_ = 5 × *k*_*b*2_ (green color) keeping all other rates constant same as in Table S13 **C)** For case **T**_3_, the model simulated time profiles of **CI)** product (*X*) and **CII)** product (*Y*), when [C-D] events (4 [C-D]) fine-tune the intermediate rate *k*_i1_ and the catalytic rate *k*_*x*_ collectively (solid lines). Time profiles of **CIII)** product (*X*) and **CIV)** product (*Y*), when [C-D] event (4 [C-D]) alters the intermediate rate *k*_ii1_ and the catalytic rate *k*_*y*_ collectively (solid lines) compared to CT. The simulation has been performed by increasing the binding rate as *k*_*b*2_ = 2 × *k*_*b*1_(cyan color) and *k*_*b*2_ = 5 × *k*_*b*1_(green color) keeping all other rates constant same as in Table S13. The dotted lines represent the simulated curves for CT (Table S13). The dashed lines represent the product formation for the increment of bimolecular rate (*k*_*b*1_/*L*, *k*_*b*2_/*L*, *L* = 0.1) due to the compression of length (*L*), while performing 4 [C-D] cycles.

For *C*_2_ case, if the [C-D] cycles increase the reaction flux via influencing the catalytic rate *k*_*x*_ along with decreasing the length of the material transiently, the production of *X* enhances 2 folds (Fig. 3CI, S9BI) compared to CT. However, it decreases the amount of *Y* significantly (solid lines, Fig. 3CII, S9BII), if only the length of the material gets influenced due to the [C-D] cycles (dotted lines, Fig. S9BII). Though, the system has a moderately higher propensity to form the product *X* (red lines, Fig. 3AIII), still, it is possible to increase the production rates of *Y* almost 3 folds (Fig. 3CIV, S9BIV) without almost altering *X* (Fig. 3CIII, S9BIII) compared to CT, if [C-D] cycles happen to alter the transition rates (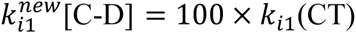). In contrast, for case *C*_3_, altering *k*_*x*_ or *k*_i1_ leads to a similar analysis as observed for scheme-II. These observations can be summarized through a probable energy profile diagram (Fig. S10) under different conditions, which suggests that by fine-tuning the reactions via [C-D] cycles, the propensity of product formation can be altered systematically.

**Fig. 2.**
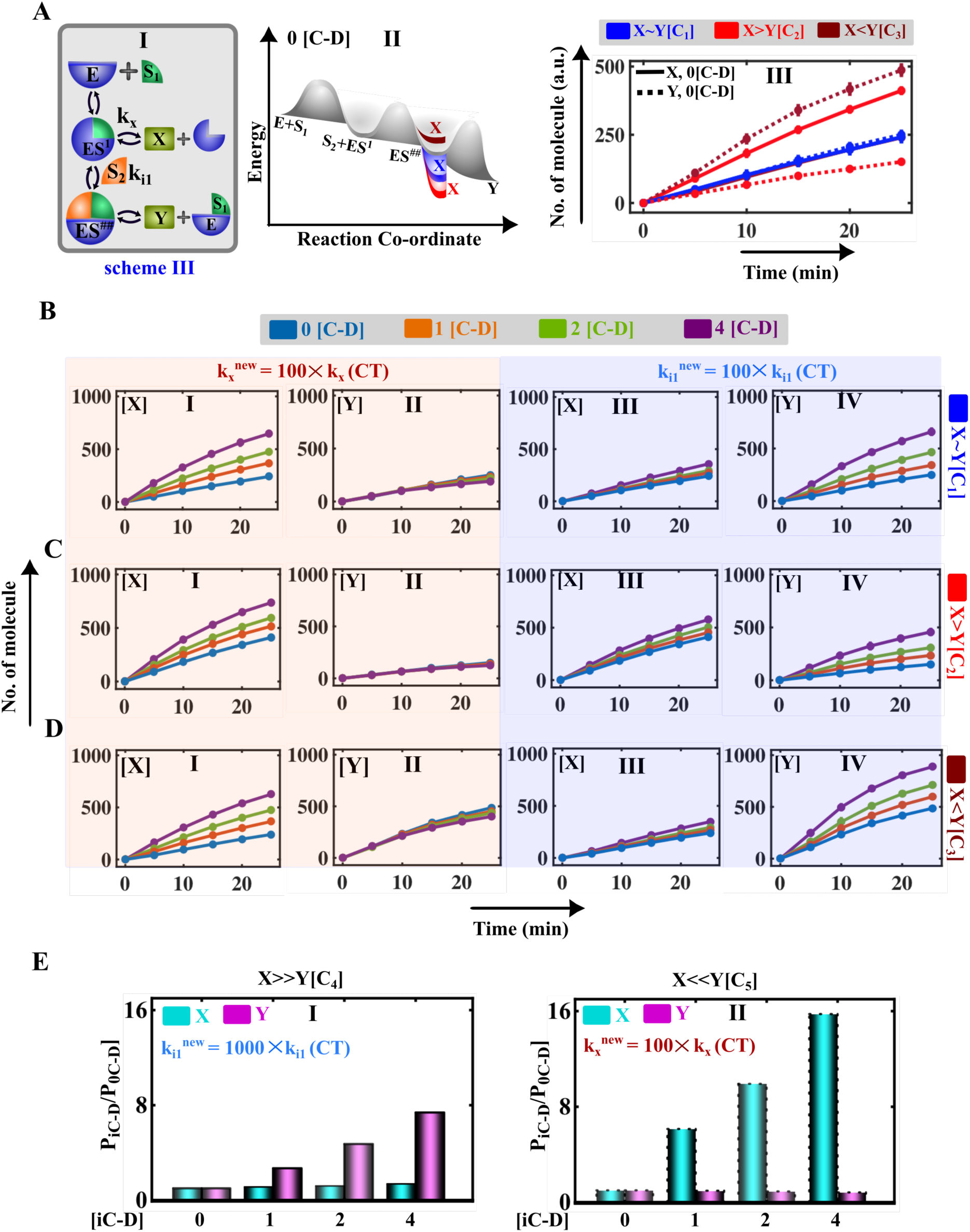
Kinetic study of the substrate transduced product formation (scheme-III) in the absence and presence of the [C-D] cycles. **A) (I)** Scheme-III describes an enzymatic reaction, where a substrate (*S*_1_) reacts to an enzyme *E* and gives the first enzyme-substrate complex *ES*^1^, which binds with another substrate (*S*_2_) and directly form a hetero-trimeric complex *ES*^##^. The enzyme-substrate complex *ES*^1^ produces the product *X* and *ES*^##^gives *Y*. **(II)** Probable free energy profile diagram for the catalytic reaction following scheme-III under *C*_1_, *C*_2_, and *C*_3_ conditions.^50^ Time course simulations of the **(AIII)** product (*X*), and **(AIV)** product (*Y*) under three different scenarios (*X*∼*Y*, *C*_1_), (*X* > *Y*, *C*_2_) and (*X* < *Y*, *C*_3_), in the absence of [C-D] cycles (corresponding parameter set is provided in Table S9). The amount of product *X* (**BI** for case *C*_1_, **CI** for case *C*_2_, and **DI** for case *C*_3_) and product *Y* (**BII** for *C*_1_ case, **CII** for *C*_2_ case, and **DII** for *C*_3_ case), if the [C-D] cycles accelerate the catalytic rate *k*_*x*_100 times compared to CT. The amount of product *X* (**BIII** for case *C*_1_, **CIII** for case *C*_2_, and **DIII** for case *C*_3_) and product *Y* (**BIV** for *C*_1_ case, **CIV** for *C*_2_ case, and **DIV** for *C*_3_ case), when the [C-D] cycles fine-tune the conversion rate as (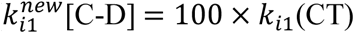). The product formation for scheme-II depicts two extreme scenarios, where *X* >> (solid lines, *C*_4_) and *X* << *Y*(dotted lines, *C*_5_), without any compression-decompression experiment (0 [C-D] cycle) using the parameter set provided in (Table S9). **(E)** The relative increment of the level of products *X* and *Y* (where *P*_i*CD*_/*P*_0*C*D_, *i* = 1, 2, 4) by altering the **(I)** *k_x_* (1000 times) and **(II)** *k*_*x*_(100 times) under different [C-D] cycles for the cases *C*_4_ and *C*_5_, respectively.

Similar to scheme-II, scheme-III has been analyzed for two extreme conditions. (1) The formation rate of *X* (*k*_*x*_) is substantially higher and the binding rate (*k*_i1_) of the substrate *S*_2_ with *ES*^1^ is very weak (Case-*C*_4_, Table S9). At this condition, the propensity of forming *X* is much higher than *Y* (Fig. S11A). Under this situation, elevating the conversion rate (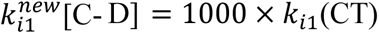) by even 1000 folds results in marginal increment in *Y* (Fig. 4b.3EI, S11(C-D)). (2) On the other hand, the stronger binding affinity of the substrate *S*_2_ with *ES*^1^ and poor catalytic rate for *X*, *k*_*x*_(Table S9), significantly tilts the system towards the accumulation of *Y* (Fig. S11B). At this state, if the [C-D] cycles have the ability to elevate the catalytic rate by at least about 100 folds, even then it increases the production of *X* almost six folds by reducing the formation of *Y* (Fig. 3EII, S11(E-F)).

The observation based on scheme-II and scheme-III reveals that the creation of *X* is dependent on the first enzyme-substrate complex (*ES*^1^, *ES*). Further, the simulation analysis performed on scheme-III depicts that alteration of the transition rate between the intermediate conformations (*ES*^1^ to *ES*^##^) can significantly modulate the production of *Y*. As, for both of the cases, *X* accumulates at a faster time scale than as compared to *Y*, by applying [C-D] cycles the reaction propensity can be easily directed toward this pathway beyond wherever the system exists.

### Dynamical switching of the kinetics for the competitive binding of the substrates encountering [C-D] cycles (scheme-IV)

The previous analysis illustrates the dynamic role played by the two different substrates, where one substrate binds directly with the enzyme and the other substrate binds directly with the enzyme-substrate intermediate conformers. Often, the presence of two substrates encountering competitive binding for the enzyme has been observed in many biological systems.^51–54^ Thus, a schematic has been designed (Fig. 4A, S1D and Table S(10-11)) inspired by the ping pong bi bi mechanism with an additional modification to reveal the relative product formation abilities from two different intermediate conformers under [C-D] cycles. Here, two substrates (*S*_1_ and *S*_2_) can bind with a single enzyme (*E*) creating two distinct enzyme-substrate complexes *ES*^1^ and *ES*^2^, which can initiate cascade reactions by further reacting with either substrates *S*_2_ (with *ES*^1^) or *S*_1_ (with *ES*^2^) to produce another intermediate state *ES*^##^. These intermediate states can reversibly interconvert with each other. The intermediate *ES*^1^ generates the product *X* and the enzyme *E*, whereas *ES*^##^converts to product *Y* and the conformer *ES*^1^.

The relative levels of *X* and *Y* will be dynamically controlled and modulated by the competitive binding affinity of the substrates with the enzymes. A probable free energy diagram of the mechanism has been depicted in Fig. 4B, where the kinetics of the product formation has been outlined with various competitive binding probabilities (**T**_1_, **T**_2_ and **T**_3_, Table S11). The length contraction effect due to the [C-D] cycles (alter the second order bimolecular reaction rates (*k*_*b*1_, *k*_*b*2_)) doesn’t significantly modify the product formation rate appreciably (Fig. S12) for scheme-IV. At this point, it is possible to consider three different situations that may arise from the competitive substrates binding affinity with the enzyme for scheme-IV (Fig. 4B). (1) The two substrates can have almost equal probability (*k*_*b*1_ ≈ *k*_*b*2_) to bind with the enzyme (gray shades (**T**_1_ case) in Fig. 4B). In this case, if the [C-D] cycle (4 [C-D]) alters the corresponding catalytic rates (*k*_*x*_ or *k*_*y*_), then it aids up the formation of the product *X* and *Y* (solid lines in Fig. S13). (2) At a low binding rate of substrate *S*_2_, the reaction scheme follows Michaelis-Menten-like kinetics (case-**T**_2_), whereas for the very low binding affinity of the substrate *S*_1_ with enzyme (case-**T**_3_), the scheme-IV converts to almost scheme-III. Afterward, analysis has been performed for only moderate and higher binding affinity for both substrates.

However, the substantially higher binding affinity of the substrate *S*_1_ compare to the substrate *S*_2_ (*k*_*b*1_ ≈ *n* × *k*_*b*2_, *n* = 1,3,10) yields the product *X* at least 2-3-fold higher than *Y* (dashed lines in Fig. S14(A-B)). In this situation, if [C-D] cycles are able to enhance the catalytic rate *k*_*x*_ that lead to an abrupt increase in the amount of *X* (solid lines, Fig. S14A), reducing the production of Y drastically (solid lines, Fig. S14B). The formation of the product *Y* simultaneously produces the complex *ES*^1^, thus favoring the formation of product *X*. The enhancement of the catalytic rate *k*, helps to accumulate the product *Y* (solid lines, Fig. S14C), but can’t reduce the product *X* much compare to CT (Fig. S14D). Here, it is very important to note that [C-D] cycles may alter the reversible conversion rates (*k*_i1_, *k*_i2_, *k*_ii1_, *k*_ii2_) related to intermediate conformers *ES*^1^, *ES*^2^ and *ES*^##^ and lead to diverse kinetic effects on product formation. For example, if [C-D] cycles escalate the *k*_i1_, the production of *Y* increases almost six folds without changing the *X* level (solid lines, Fig. 4C(I-II)) compare to CT. However, altering *k*_ii1_ rate doesn’t accelerate the production rate of either *X* or *Y* in a significant manner (solid lines, Fig. 4C(III-IV)). Interestingly, accelerating the intermediate rate *k*_i2_ or *k*_ii2_ have a marginal effect on the production rate (Fig. S15). (3) For case **T**_3_, when the substrate *S*_2_ has higher binding affinity than the substrate *S*_1_ (*k*_*b*2_(*C***T**) ≈ *n* × *k*_*b*1_(*C***T**), *n* = 1,3,10, **T**_3_), changing the catalytic rate either *k*_*x*_ or *k*_*y*_, the production of *X* shows an abrupt enhancement and the yield of *X* increases at least 2-3-fold (Fig. S16). At the same time, increase in the *k*_i1_ upon [C-D] cycles results in the moderate enhancement of *Y* with little reduction of *X* (solid lines, Fig. 4D(I-II), S17(A-B)). An interesting observation has been found that the alteration of *k*_ii1_ under [C-D] cycles not only increase the amount of product *Y* but also accelerate the production of *X* (solid lines, Fig. 4D(III-IV), S17(C-D)). This important observation reveals that even under low binding affinity of the substrate *S*_1_, there are possibilities to enhance the accumulation of product *X* if the enzymatic reactions of biomaterials following the kinetic pathway like scheme-IV undergo [C-D] cycles.

The scheme-IV exhibits a complex mechanism, where particular substrates are responsible to give a particular product (e.g., substrate *S*_1_ gives product *X* and substrate *S*_2_ yields *Y*). Also, their productions are dependent and connected with each other via trimeric intermediate conformers *ES*^##^. For this kind of system, the abovementioned analysis finds that by employing [C-D] cycles on biomaterials, it is possible to elevate the production in many ways, which is generally not attainable for reactions occurring in normal systems (not within biomaterials). Even sometimes, it is possible to enhance both of the products by only modulating one particular catalytic rate through altering [C-D] cycles.

### Kinetic study of the competitive binding of the enzyme to substrate undergoing [C-D] event (scheme-V)

In biological systems, often the competitive binding affinity of various enzymes for a particular substrate has been observed.^55,56^ Thus, a two-enzyme and one-substrate system within these biomaterials have been designed (schematic-V) and a kinetic study has been performed using the established computational methodology. Here, two enzymes (*E*_1_ and *E*_2_) can undergo competitive binding for the same substrate (*S*) based on the different binding affinity of the enzyme with the substrate (Fig. 5A, Table S12). (1) **T**_1_, two enzymes have an almost similar binding affinity ([*k*_*b*1_ ∼ *k*_*b*2_], Fig. 5AII) with *S*. It has been found that in the case of **T**_1_, alterations made in only the catalytic rate (*k*_*x*_, *k*_*y*_) have very less effect on the product formation rate (Fig. S18(A-D), S19(A-D)) compared to changing the conversion rate (*k*_i1_, *k*_ii1_) under the chosen parametric conditions (Table S13). The enhancement of the conversion rate (*k*_i1_, *k*_ii1_) along with catalytic rate (*k*_*x*_, *k*_*y*_) during [C-D] cycles increase the formation of either product *X* or *Y* consecutively compare to CT (Fig. S18(E-F), S19(E-F)). (2) For case **T**_2_, enzyme *E*_1_ have higher binding affinity with *S* than *E*_2_ ([*k*_*b*1_ > *k*_*b*2_], Fig. 5AII). In this case, if the [C-D] cycles can fine-tune the reaction propensity towards the formation of *X* (influencing *k*_i1_ and *k*_*x*_), the product (*X*) formation get enhanced, consequently decreasing the production of *Y* compared to CT (solid lines, Fig. 5B(I-II)). For higher values of *k*_*b*1_, an abrupt increase in the production of *X* has been found, which saturates with time and results in an almost four-fold reduction in *Y* (solid green lines, 5B(I-II)). On the contrary, enhancing all the reaction fluxes related to *Y* (conversion rate *k*_ii1_ and catalytic rate *k*_*y*_), moderately changes the conversion of *Y* compared to CT (Fig. 5B(III-IV)). (3) In case of **T**_3_, enzyme *E*_2_ have a higher binding affinity with *S* than *E*_1_ ([*k*_*b*2_ > *k*_*b*1_], Fig. 5CII), similar results have been observed as found for **T**_2_ on product formation rate by performing similar analysis altering the catalytic rate along with conversion rate (*k*_*x*_/*k*_i1_ and *k*_*y*_/*k*_ii1_, Fig. 5C).

**Fig. 5.**
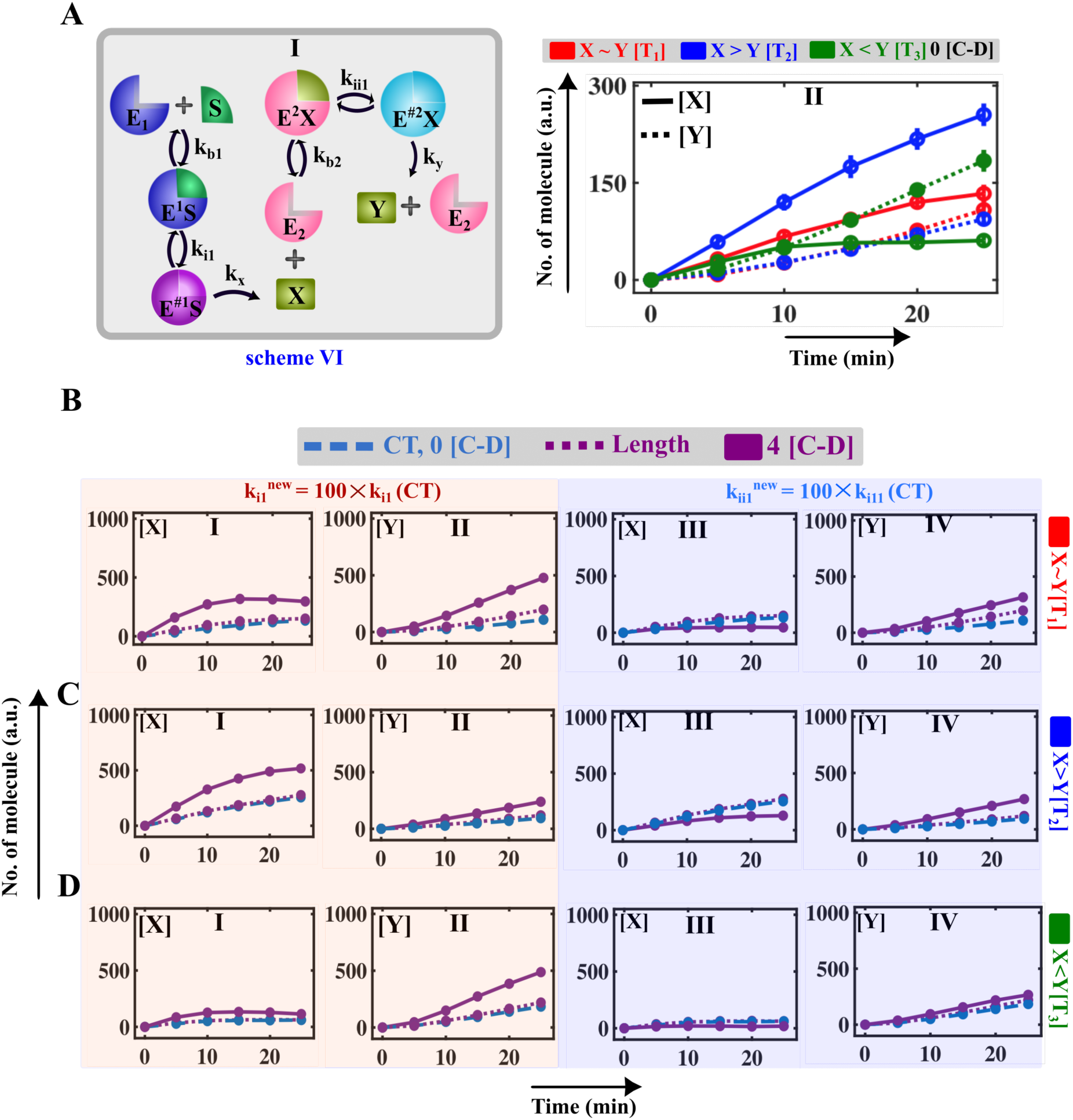
Fine-tuning of the formation of products **X** and **Y** produced from biochemical reaction following scheme-VI due to [C-D] cycles. **AI)** The proposed mechanism consists of two enzymes (*E*_1_ and *E*_2_) having the substrate-specific binding capability where the *E*_1_ can only bind with substrate *S* yielding an intermediate conformer *E*^1^*S*, which eventually converts to another conformer *E*^#1^*S*. The intermediate *E*^#1^*S* dissociates into product *X* and *E*_1_, and *X* acts as the substrate for another enzyme, *E*_2_ by similarly forming two sequential intermediates *E*^2^*X* and *E*^#2^*X* and end up in forming another product *Y* along with regenerating the enzyme *E*_2_. **AII)** Time course simulations of the product (*X*) (solid lines) and product (*Y*) (dotted lines) under three different scenarios (Table S13) (*X*∼*Y*, **T**_1_), (*X* > *Y*, **T**_2_) and (*X* < *Y*, **T**_3_) in the absence of [C-D] cycles (corresponding parameter set is provided in (Table S13)). For case **T**_1_, **(B)**, case **T**_2_, **(C)** and case **T**_3_, **(D)**, the simulated time profiles of **I)** product (*X*) and **II)** product (*Y*), when [C-D] event (4 [C-D]) fine-tunes the intermediate rate *k*_i1_ and the time evolution of **III)** product (*X*) and **IV)** product (*Y*), when [C-D] event (4 [C-D]) alters the intermediate rate *k*_ii1_ compared to CT. The dotted lines represent the simulated curves when no rate has been altered (CT, Table S13. The dashed lines represent the product formation for the increment of bimolecular rate (*k*_*b*1_/*L*, *k*_*b*2_/*L*, *L* = 0.1) due to the compression of length (*L*), while performing 4 [C-D] cycles keeping all other rates constant same as in Table S13.

### The kinetics of an enzyme-catalyzed reaction where the *in-situ* product is used as substrate (scheme-VI)

Using my proposed computational method, I have been able to predict particular strategies to improve product formation for various enzyme-catalyzed reactions which will help in designing new experimentally testable hypotheses. For example, the proposed methodology can be applied to understand heterogeneous catalysis following almost a Ping-Pong Bi Bi kind of mechanism, where the in-situ product of the first enzyme-substrate complex has been used as the substrate for the second enzyme (Fig. 6AI, S20). In order to search particular effective kinetic rate constants involved in scheme VI, again I consider three distinct cases (Fig. 6AII, S21, Table S(14-15); (i) **T**_1_, both the product reproduced in almost comparable amounts, (ii) **T**_2_, the formation rate of product *X* is significantly higher than the product *Y* and (iii) **T**_3_, production of *Y* is higher than the creation of *X*. I assume that all probable reaction fluxes during [C-D] cycle implementation may get altered under different circumstances. As [C-D] cycles may alter any reaction fluxes associated with scheme-VI depending on reaction conditions and the structure of the material. Here, all such possibilities have been explored by fine-tuning the kinetic rates. It has been found that changes made in any catalytic rates (*k*_*x*_, *k*_*y*_) don’t accelerate the accumulation of *X* and *Y* significantly (Fig. S(22-23)) under all parametric conditions (Table S15). However, due to the specific nature of the mechanism, the reaction flux associated with the conversion rate (*k*_i1_, *k*_i2_) has a significant effect over the catalytic rate. As the formation of *X* facilitates the creation of *Y*, increasing the transition rate *k*_i1_ enhances the production of both products under all conditions (Fig. 6B(I-II), C(I-II), and D(I-II)). In contrast, increasing the reaction rate *k*_ii1_ not able to enhance the production rate of *Y* and *X* (Fig. 6B(III-IV), C(III-IV), and D(III-IV)), even when the system is significantly tilted towards the formation of *Y*(for case **T**_3_).

**Fig. 6.**
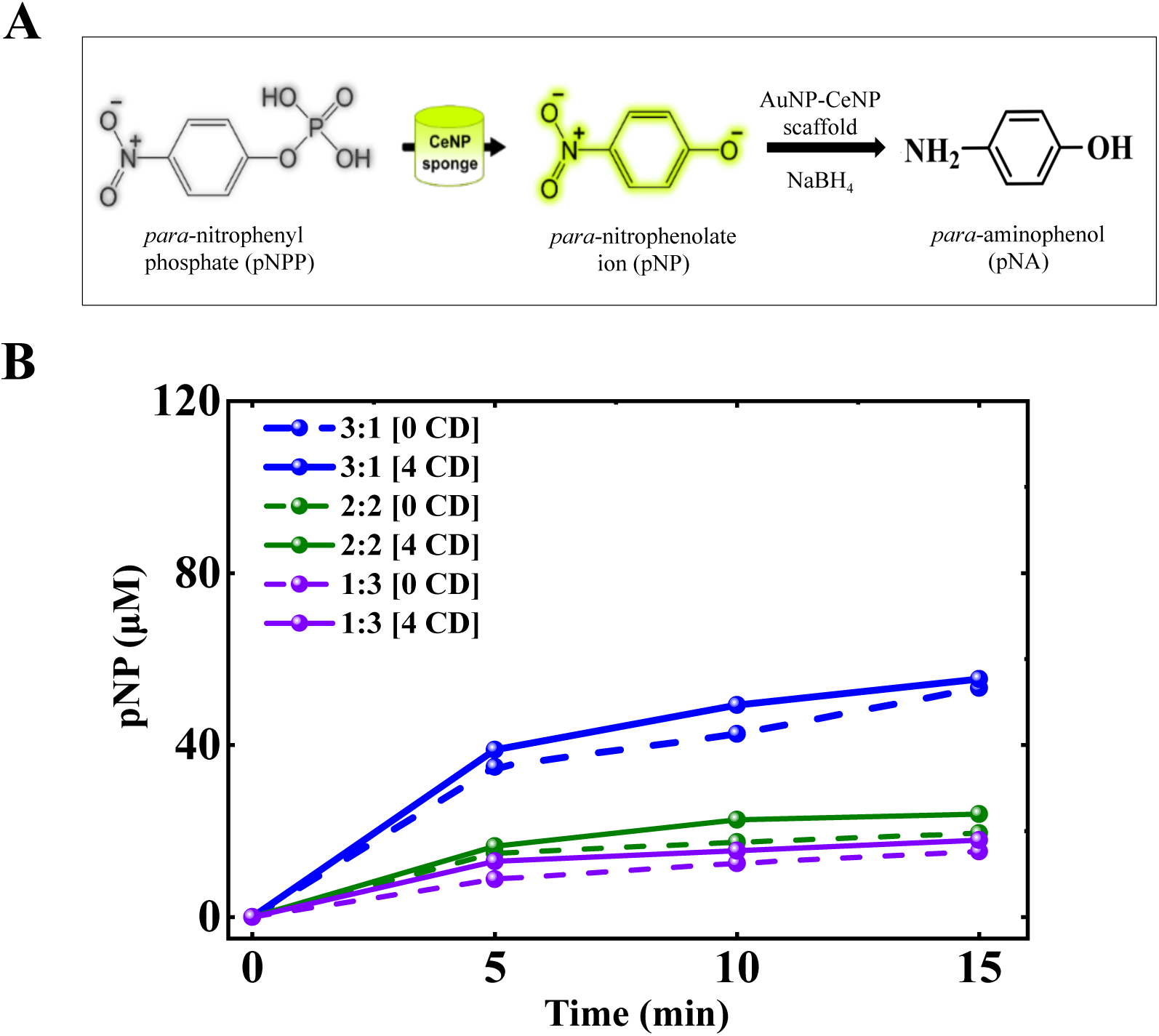
Heterogeneous catalysis of the consecutive substrate conversion reaction in a confined environment. **A)** Hydrolysis of para-nitrophenyl phosphate (pNPP) into para-nitrophenylate (pNP) by ceria nanoparticle (CeNP). In the presence of NaBH_4_, gold nanoparticles (AuNP) reduced the pNP to PNA. **B)** Hybrid scaffold has been prepared by Lohitha *et al.* combining the CeNP and AuNP in different ratios ((1) 3:1 (CeNP = 2250 μM, AuNP = 750 μM), (2) 2:2 (CeNP = 1500 μM, AuNP = 1500 μM), (3) 1:3 (CeNP = 750μM, AuNP = 2250μM). Comparison of the time evolution of product pNP for 0 [C-D] and undergoing 4 [C-D] cycles for all variations of combined nanoparticles.

Experimentally, it is possible to design such a system (scheme-VI), where heterogeneous catalysis has been observed. Hegde *et al.* has synthesized the ceria nano-particle-based biomimicking materials that can hydrolyze pNPP.^14^ By collaborating with Prof. Kamendra P. Sharma’s Lab, Hegde *et al.* has synthesized a ceria and gold nano-particle-based scaffold (CeNP & AuNP) and tried to observe the enzymatic reaction for two enzymes system. CeNP hydrolyses the substrate pNPP into pNP, and AuNP converts this in-situ product pNP into pNA in the presence of NaBH_4_ (Fig. S20). The evolution of pNP has been observed for three different ratios of CeNP and AuNP under the [C-D] cycle (Fig. 7).

### Summary

The analysis of the kinetics of different schemes under [C-D] cycles sheds light on the importance of using theoretical and computational methodologies in understanding the biochemical reaction happening within a small scaffold. My proposed methodology helped in analysing these kinetic models and predicts ways to optimize such complex biocatalytic reactions happening within these spongy materials. In literature, mostly three different deterministic approaches have been found to study the kinetics of enzymatic reactions: transition state, steady state, and rapid equilibrium kinetics.^57,58^ The transition state theory has been applied for rapid reactions and is linked with mainly structures of the enzyme.^57^ The steady state concept has been used for the system that reaches a steady state quickly.^57^ The rapid equilibrium kinetics methods are used to analyze reaction systems where the species involved in the reaction are mostly in equilibrium with each other.^58,59^ The MM expression has been extensively used to understand how the rate of the reaction depends on the substrate concentration. ^51,52,60,61^

However, fluctuations of both quantum mechanical and thermal origin are inherently embedded into the natural systems.^62,63^ At the molecular level, this noise affects the turnover rate of the single-molecule enzyme-substrate reactions.^38,39,41,43,64^ In addition, for inhomogeneous systems, diffusion may play a significant role in controlling the catalytic output.^38,65–67^ The intrinsic spatial and temporal fluctuations of the catalytic rate of such reaction-diffusion systems can be statistically measured by applying stochastic formulations.^41,45,68–72^ Numerous methods have been developed to improve the efficiency of such hybrid computational methods.^73–75^ However, it becomes extremely challenging and very few methodologies have been developed to study the reaction of such reaction-diffusion systems occurring within a biomaterial undergoing mechanical stress. Here, I have proposed a computational approach that can easily reveal interesting observations about the kinetic progression of complicated enzyme-catalyzed reactions within such biomaterials. Thus, it provides a way to circumvent the difficulties that one might experience while performing these reactions in the future within such biocatalytic materials of any dimension. Hence, my computational studies will help to reduce the time and cost associated with performing these reactions in these spongy materials for various application purposes, which is highly relevant for industrial production. From a catalysis perspective, my proposed method can suggest new strategies to rationally design such biomaterial that can enhance the production of the complex biochemical system exhibiting such mechanoresponsive property.

## Notes

### Competing Interest Statement

The authors have declared no competing interest.

