## Supplementray for "Understanding the dynamics of complex *in-situ* enzyme-catalyzed reactions undergoing mechanical stress"

### Text S1 Theoretical methods

#### A) Proposition of a modified stochastic methodology of reaction diffusion system.

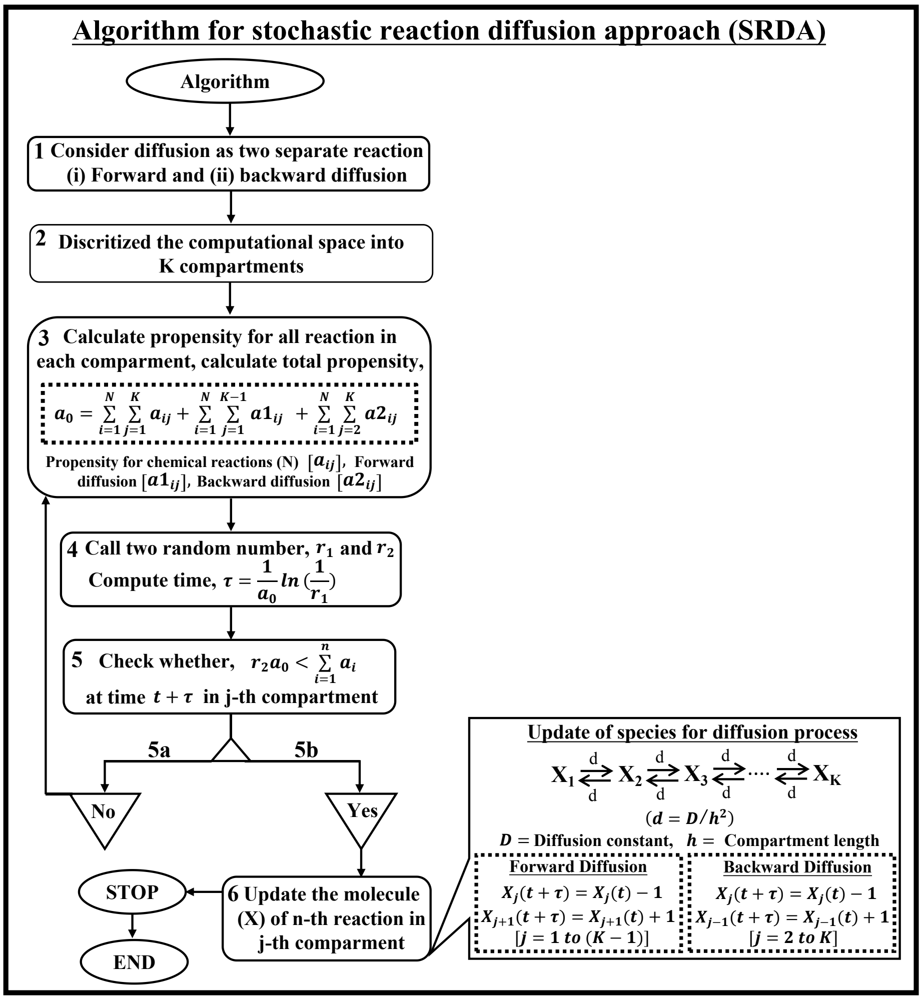

**Module S1.** Algorithm for implementing modified stochastic reaction diffusion approach (SRDA).

A stochastic description is needed to explain the spatiotemporal fluctuation in the concentration of chemically reacting species, which mainly arises both from diffusion and reaction processes. The fluctuation and randomness in the chemical reactions can be formulated by using Gillespie's stochastic simulation algorithm (SSA) (ref). However, this assumption gets violated where the number of molecule is very low and the system exhibit a spatio-temporal inhomogeneity along with the stochasticity which is often observed inside the living cells. Additionally, the spatial inhomogeneity may also arises for the porous structure of a biomaterial. One of the ways to simulate this type of system is to partition the space into small homogenous spatial domains by assuming that the stochastic events can only occur within the same compartment and molecule can diffuse to its neighboring compartments.

The fluctuation and randomness in the chemical reactions can be formulated by using Gillespie's stochastic simulation algorithm (SSA). However, SSA hypothesis can be applied to capture diffusion mediated heterogeinity in the reaction system. Erban *et al.* have been provided a methodology to compute the time evolution of any molecule encountering diffusion in compartmentalized system by considering diffusion as a separate chemical reaction using SSA. However, their (SSA) based computational method formulates the update of the species in such a way that the calculation of propensity step repetitively appear after occurring each elementary process thereby lengthening the existing numerical method. We have modified their approach by separating the diffusion as forward and backward diffusion **(Step-(1-2) of Module 1)** and have calculated the propensity functions separately for this two processes **(Step-(3) of Module 1)**. Afterwards, two random numbers have been used for finding the probability of occurring n-th reactions in between time $t$ and $(t + \tau)$ **(Step-(4-6) of Module 1)**. The division of diffusion into two distinct processes allowed to update all the reaction and diffusion events within a closed loop **(Step-5a,5b and 6 of Module 1)**. Thus, our proposed methodology helps to simulate the stochastic reaction diffusion process with better efficiency and reduces the computational time.

**B) Numerical method to incorporate the compression-decompression phenomena in SRDA algorithm (NMCD)**

Consequently, we have developed a generalized numerical recipe to include external effect on enzymatic reaction. Here, we focused to implement the [CD] event numerically to mimic the effect of compression-decompression experiment [CD] on enzymatic reaction occurring within a biomaterial performed by Mehak et *al*. To validate their enzymatic reaction performed using spongy biomaterial, we introduced two separate time scales in our simulation to indicate (i) the time span for the normal chemical reactions $(t_{f1})$ and (ii) the time duration for performing the mechanical operations $(t_{f2})$ due to CD cycle. The $(t_{f2})$ helps to define the different frequency of [CD] cycles (for (1 CD) $\left(t_{f2} = 1\right)$, for (2 CD) $\left(t_{f2} = 0.5\right)$ and for (4 CD) $\left(t_{f2} = 0.25\right)$). The algorithm start by defining these two characteristic time scales and setting the initial time, $t_i = 0$ (with a counter $Icount(= 0)$) **(Step-(1-2) of Module 2)**. After that, we allowed the system to evolve following the SRDA protocol for either $t_{f1}$ or $t_{f2}$ and continue to track the total of every species involved in the enzymatic reaction after completing each iteration **(Step-(5-6) of Module 2)** and performed simulation for desire reaction duration under varied [CD] cycle conditions **(Step-9 of Module 2)**.

**C) Incorporation of the effect of the alteration of length on kinetics due to [CD] phenomena in existing (NMCD) method**

From the experiments, it is evident that the compression activity leads to the mixing of the molecules by discarding about 60% of the substrate and product molecules from the spongy materials. Whereas, the decompression process brings back all the substances into the system again by redistributing them into different compartment. We implemented this idea into our algorithm by calculating the total amount of the substrate and product in similar proportions separately **(Step-(7-8) of Module 2)** and redistributed the molecule **(Step-(5-5a) of Module 2)** after each [C-D] cycle inside the compartments. To distribute the reacting species (enzyme for only $Icount(= 0)$, substrate after each $Icount(= 0,1, \ldots.)$ and product after each $Icount(= 1,2, \ldots.)$), we have used either the Box-Mullar transformation or binomial distribution, by carefully choosing them based on the total amount of molecule presents in the system. The counter ($Icount$) has been updated after each distribution (**Step-(7-8) of Module 2)** and formulated in such a way that it appropriately distributes the molecule (substrate and product) after each [C-D] cycle **(Step-5a of Module 2)**.

**D) Addition of the alteration of kinetic rates during [CD] cycle in existing (NMCD) method**

As only incorporation of the [CD] cycle or length factor was not sufficient to explain the experimentally observed enhancement of the product formation, we included the characteristic feature, if the [CD] cycle could able to enhance the catalytic efficiency by altering the reaction rates (catalytic rate or interconvertible rate). In module 5b, we have included this concept numerically considering rising reaction rate with altering [CD] cycle **(Step-(5-5b) of Module 2)**. To explain the enhancement of product under increasing frequency of [C-D] cycle, we have assumed that the particular reaction rates (either catalytic rate or conformations transition rate) get enhanced during the time span for compression and decompression cycle **(Step-5a of Module 2)** into the existing method. The simulation ends after a certain time (in our case at 25 minutes), which has been incorporated by introducing the sub-module 8a.

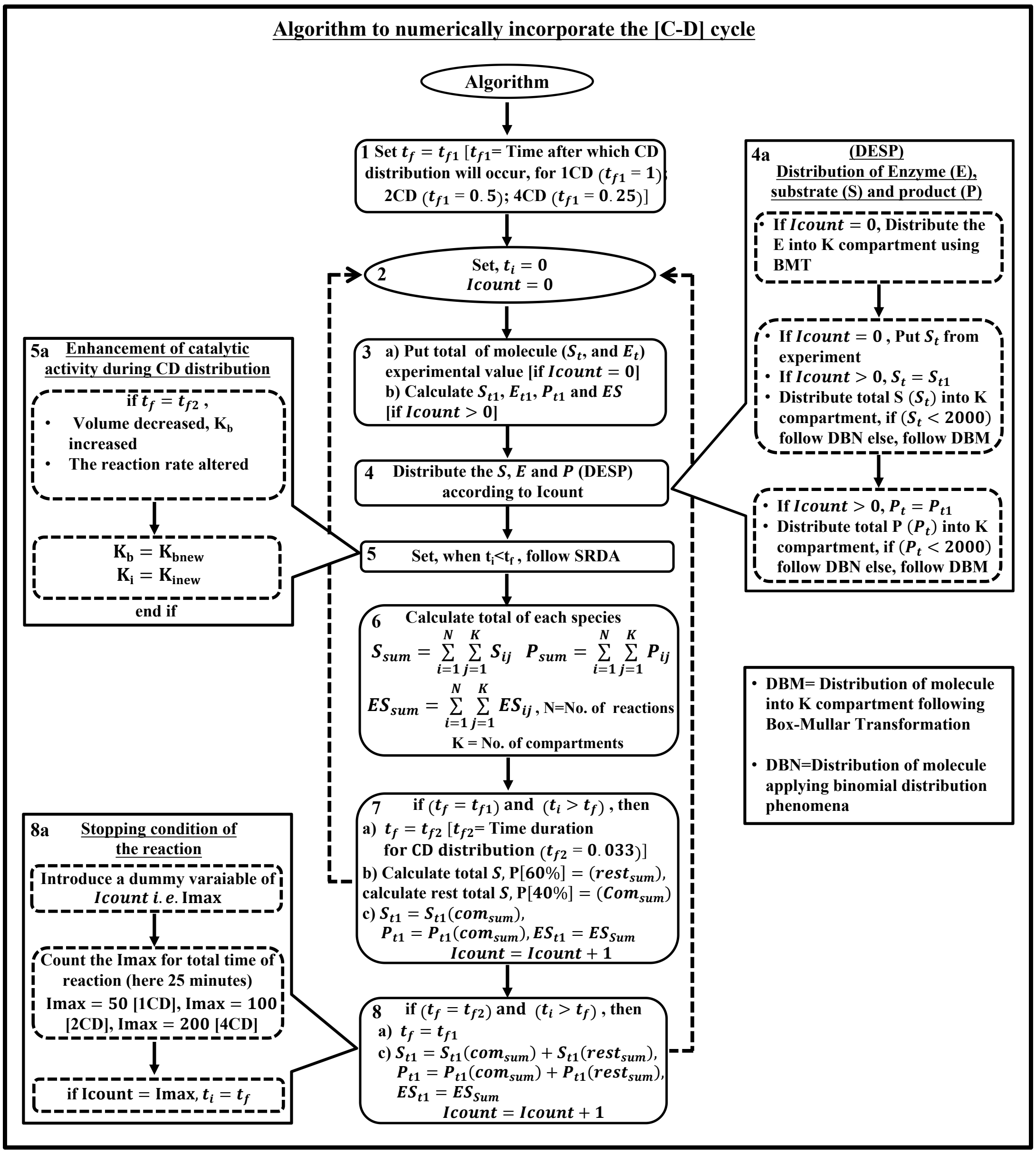

**Module S2.** Algorithm to incorporate the [C-D] phenomena into existing SRDA method.

particular time **(Step-4a of Module 2).**

#### Text S2 Probable schematic for enzyme-catalyzed reaction.

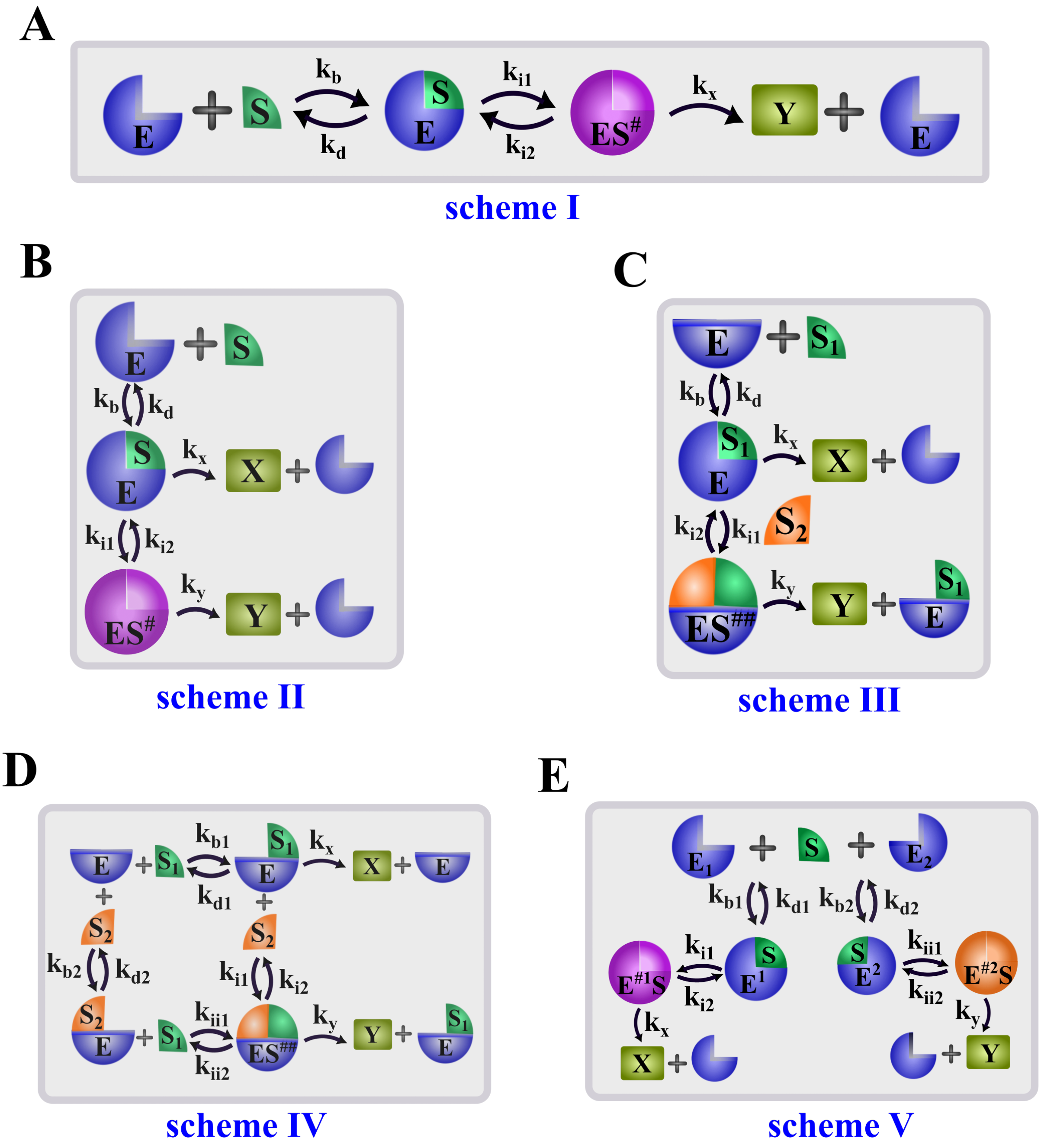

**Fig. S1 Schematic representation of different probable mechanisms of enzymatic reaction occurring within a spongy bio-material.**

The mechanisms can proceed involving multiple substrates ($S_1$ and $S_2$), multiple enzymes ($E_1$ and $E_2$), multiple interconvertible states ($ES, ES^{\#}$, and $ES^{\#\#}$), which can produce single or multiple products ($X$ and $Y$).

#### Text S3 Calculations:

##### A) Calculation of the porous compartment present inside the biomaterial

The total length of the biomaterial $= 1\ cm = 10\ mm = 10{,}000\ micron$
Size of each pore $\approx\ 200\ micron$
Total number of the compartment $\approx\ (10{,}000/200)\ \approx\ 50$

##### B) Calculation of the enzyme-substrate ratio

Weight of the substrate (pNPP) taken $= 0.000556725\ g$
Number of moles of pNPP $= (0.000556725/371.15) = 0.15 \times 10^{-5}$ moles
The number of the molecule of pNPP present in the sponge of size $(1\ cm\ \times 1cm) = 0.15 \times 10^{-5}\ \times\ 6.023\ \times\ 10^{23}$
Weight of the enzyme (ALPs) taken $= 0.2 \times 10^{-3}\ g$
The number of moles of enzyme $= (0.2 \times 10^{-3}/86000) = 0.232 \times 10^{-8}$ moles
The number of the molecule of enzyme present in the sponge of size $(1\ cm\ \times 1cm) = 0.232 \times 10^{-8}\ \times\ 6.023\ \times\ 10^{23}$
The ratio of the molecule of the substrate to the enzyme is $\sim\ 650{:}\,1$

##### C) Converting the diffusion constant into 1-dimension

Length of each compartment (h) = (Total length of material/total number of the compartment) $= \left(\frac{1\ cm}{100}\right) = 0.01\ cm$
Diffusion constant per unit length = $\left(\frac{D_h}{h^2}\right)$, $D_h$ = Diffusion constant of the material

##### D) Alteration of the length into 1-dimension

It is already well-known that, $\left(c = \frac{n}{V}\right)$, where C= Concentration of the species, n=Number of mole =n0/NA, no=Number of molecules, NA= Avogadro number, V=volume. Thus, the second order rate constant ($k_b$) is inversely proportional to the volume, due to the contraction of the spongy material by compression, the volume gets reduced which increased the bimolecular rate $k_b$.

For a 1-dimensional system, the volume is equivalent to length. During the compression experiment the $k_b$ will be changed and updated as $\left( k_{bnew} = \frac{old\ k_b}{length\ changed} \right) = \left( new\ k_b = \frac{old\ k_b}{L} \right)$.

#### Text S4 Observation of the kinetics of the biochemical reaction following mechanisms like scheme-I

##### A) ODEs of the proposed model

$$\frac{dES}{dt} = (k_b \times E \times S) - (k_d \times ES) - (k_{i1} \times ES) + (k_{i2} \times ES^{\#}) \qquad [S1.1]$$

$$\frac{dES^{\#}}{dt} = (k_{i1} \times ES) - (k_{i2} \times ES^{\#}) - (k_x \times ES^{\#}) \qquad [S1.2]$$

$$\frac{dY}{dt} = (k_x \times ES^{\#}) \qquad [S1.3]$$

##### B) Alteration of the rate in scheme-I during [C-D] cycles

I have introduced the following phenomenological expression $[A5.1]$ for the enhancement in the interconvertible rate. Once this expression for the transient increase in $k_{i1}$ is introduced in the methodology, the amount of product formation with increasing [C-D] cycles obtained from simulation nicely corroborated with the experimental observation.

$$k_{i1\_new} = [(f_1 \times k_{i1}) + (f_2 \times k_{i1} \times index)] \qquad [S1.4]$$

where, $k_{i1}$ = the conversion rate $(k_{i1})$ from $ES$ to $E_1S$, $f_1 = 80$ and $b = 500, index = (1/imax), imax = 50, 100\ and\ 200$ for 1 [C-D], 2 [C-D] and 4 [C-D], respectively.

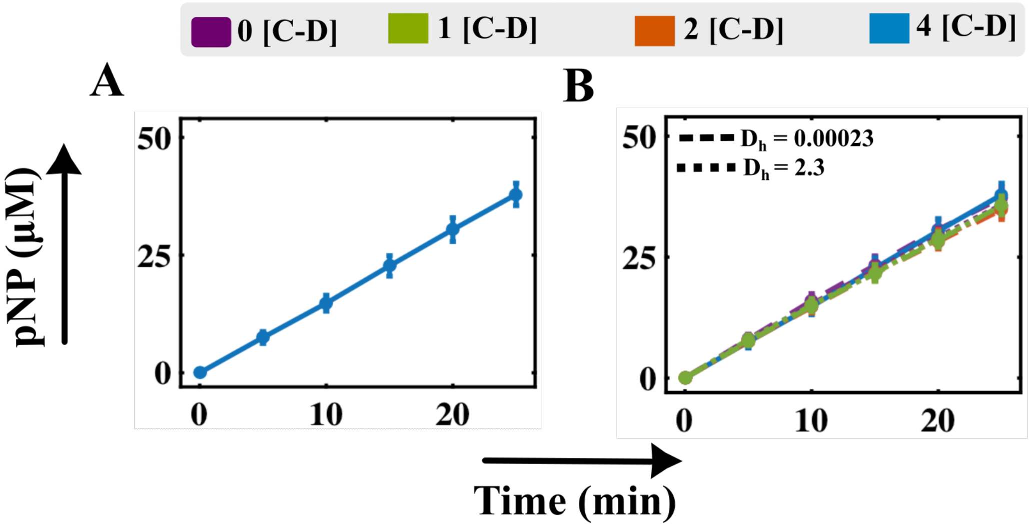

**Fig. S2 Understanding the product formation of reaction following scheme-I within a biomimetic scaffold undergoing [C-D] cycles.**

**A)** Time-course simulation of the rate of product formation by performing the SRDA methodology to reproduce the experimental data obtained by Mehek *et al.* under the 0 [C-D] cycle (CT). The kinetic rates are provided in Table 4b.2. **B)** Simulated trajectories obtained by performing NMCD simulation in case of 0, 1, 2, and 4 [C-D] cycles along with varying diffusion constant of the substrate and product $D_{hfs}/D_{hbs}$ and $D_{hfp}/D_{hbp}$, respectively (Table 4b.2, Text A5.1C).

**Table S1 Abbreviation of different variables involved in scheme-I.**

| Symbol | Description (in Conc., $\mu M$) |
|---|---|
| $E$ | The free amount of enzyme |
| $S$ | The free amount of substrate |
| $ES$ | The enzyme-substrate complex |
| $ES^{\#}$ | Another enzyme-substrate conformational state |
| $Y$ | The amount of product |

**Table S2 Description of the parameters associated with the scheme-I.**

| **Symbol** | **Description** | **Unit** | **Value** |
|---|---|---|---|
| $k_b$ | Binding rate of the enzyme ($E$) with the substrate ($S$) | $\mu M^{-1}min^{-1}$ | 0.001 |
| $k_d$ | The dissociation rate of the enzyme-substrate complex ($ES$) | $min^{-1}$ | 10 |

| | | | |
|---|---|---|---|
| $k_{i1}$ | The conversion rate of complex $(ES)$ to $ES^{\#}$ | $\mu M^{-1}$ | 0.05 |
| $k_{i2}$ | The conversion rate of complex $(ES)$ to $ES^{\#}$ | $\mu M^{-1}$ | 1 |
| $k_x$ | Catalytic rate or formation rate of the product $(Y)$ | $\mu M^{-1}$ | 2962.5 |
| $D_{hfs}$ | The forward diffusion rate of the substrate $(S)$ | $cm^2 min^{-1}$ | 0.023 |
| $D_{hbs}$ | The backward diffusion rate of the substrate $(S)$ | $cm^2 min^{-1}$ | 0.023 |
| $D_{hfp}$ | The forward diffusion rate of the product $(Y)$ | $cm^2 min^{-1}$ | 0.023 |
| $D_{hbp}$ | The backward diffusion rate of the product $(Y)$ | $cm^2 min^{-1}$ | 0.023 |
| $S_t$ | The total amount of $(S)$ | $\mu M$ | 2000 |
| $E_t$ | The total amount of $(E)$ | $\mu M$ | 80000 |
| $K$ | Total number of compartments | - | 50 |

**Table S3 Description of all the elementary reactions for the proposed scheme-I.**

| **Number** | **Reaction** | **The propensity of the reaction in $j^{th}$ compartment** |
|---|---|---|
| 1 | $E(j) + S(j) \rightarrow ES(j)$ | $a\_nu(1,j) = k_b \times E(j) \times S(j)$ |
| 2 | $ES(j) \rightarrow E(j) + S(j)$ | $a\_nu(2,j) = k_d \times ES(j)$ |
| 3 | $ES(j) \rightarrow ES^{\#}(j)$ | $a\_nu(3,j) = k_{i1} \times ES(j)$ |
| 4 | $ES^{\#}(j) \rightarrow ES(j)$ | $a\_nu(4,j) = k_{i2} \times ES^{\#}(j)$ |
| 5 | $ES^{\#}(j) \rightarrow Y(j)$ | $a\_nu(5,j) = k_x \times ES^{\#}(j)$ |
| 6 | $S(j) \rightarrow S(j+1)$ | $a\_nu(6,j) = D_{hfs} \times S(j), j = 1\ to\ (K-1)$ |
| 7 | $S(j) \rightarrow S(j-1)$ | $a\_nu(7,j) = D_{hbs} \times S(j), j = 2\ to\ K$ |
| 8 | $Y(j) \rightarrow Y(j+1)$ | $a\_nu(8,j) = D_{hfp} \times Y(j), j = 1\ to\ (K-1)$ |
| 9 | $Y(j) \rightarrow Y(j-1)$ | $a\_nu(9,j) = D_{hbp} \times Y(j), j = 2\ to\ K$ |
| **Algebraic Relation** | | |
| $E_t(j) = E(j) + ES(j) + ES^{\#}(j)$ | | |
| $S_t(j) = S(j) + ES(j) + ES^{\#}(j) + Y(j)$ | | |

#### Text S5 Observation of the kinetics of the biochemical reaction following mechanisms like scheme-II

**Table S4 Abbreviation of different variables involved in Scheme-II.**

| Symbol | Description (in Conc., $\mu M$) |
|---|---|
| $E$ | The free amount of enzyme |
| $S$ | The free amount of substrate |
| $ES$ | The enzyme-substrate complex |
| $ES^{\#}$ | Another enzyme-substrate conformational state |
| $Y$ | The amount of product |
| $X$ | The amount of another product |

**Table S5 Description of all the elementary reactions for the proposed Scheme-II.**

| Number | Reaction | The propensity of the reaction in $j^{th}$ compartment |
|---|---|---|
| 1 | $E(j) + S(j) \rightarrow ES(j)$ | $a\_nu(1,j) = k_b \times E(j) \times S(j)$ |
| 2 | $ES(j) \rightarrow E(j) + S(j)$ | $a\_nu(2,j) = k_b \times ES(j)$ |
| 3 | $ES(j) \rightarrow ES^{\#}(j)$ | $a\_nu(3,j) = k_{i1} \times ES(j)$ |
| 4 | $ES^{\#}(j) \rightarrow ES(j)$ | $a\_nu(4,j) = k_{i2} \times ES^{\#}(j)$ |
| 5 | $ES(j) \rightarrow X(j)$ | $a\_nu(5,j) = k_x \times ES(j)$ |
| 6 | $ES^{\#}(j) \rightarrow Y(j)$ | $a\_nu(6,j) = k_y \times ES^{\#}(j)$ |
| 7 | $S(j) \rightarrow S(j+1)$ | $a\_nu(7,j) = D_{hfs} \times S(j), j = 1 \; to \; (K-1)$ |
| 8 | $S(j) \rightarrow S(j-1)$ | $a\_nu(8,j) = D_{hbs} \times S(j), j = 2 \; to \; K$ |
| 9 | $X(j) \rightarrow X(j+1)$ | $a\_nu(9,j) = D_{hfp} \times X(j), j = 1 \; to \; (K-1)$ |
| 10 | $X(j) \rightarrow X(j-1)$ | $a\_nu(10,j) = D_{hbp} \times X(j), j = 2 \; to \; K$ |
| 11 | $Y(j) \rightarrow Y(j+1)$ | $a\_nu(11,j) = D_{hfp} \times Y(j), j = 1 \; to \; (K-1)$ |
| 12 | $Y(j) \rightarrow Y(j-1)$ | $a\_nu(12,j) = D_{hbp} \times Y(j), j = 2 \; to \; K$ |
| **Algebraic Relation** | | |
| $E_t(j) = E(j) + ES(j) + ES^{\#}(j)$ | | |
| $S_t(j) = S(j) + ES(j) + ES^{\#}(j) + X(j) + Y(j)$ | | |

**Table S6 Description of the parameters associated with the scheme-II**

| Symbol | Description | Unit | $C_1$ | $C_2$ | $C_3$ | $C_4$ | $C_5$ |
|---|---|---|---|---|---|---|---|
| $k_b$ | Binding rate of the enzyme $(E)$ with the substrate $(S)$ | $\mu M^{-1}min^{-1}$ | 0.005 | 0.005 | 0.005 | 0.005 | 0.005 |
| $k_d$ | The dissociation rate of the enzyme-substrate complex $(ES)$ | $min^{-1}$ | 10 | 10 | 10 | 10 | 10 |
| $k_{i1}$ | The conversion rate of complex $(ES)$ to $(ES^{\#})$ | $\mu M^{-1}$ | 3.0 | 1.0 | 3.0 | 1.0 | 28.0 |
| $k_{i2}$ | The conversion rate of complex $(ES)$ to $(ES^{\#})$ | $\mu M^{-1}$ | 1.0 | 1.0 | 1.0 | 1.0 | 1.0 |
| $k_x$ | Catalytic rate or formation rate of the product $(X)$ from $(ES)$ | $\mu M^{-1}$ | 3.0 | 3.0 | 1.0 | 30.0 | 1.0 |
| $k_y$ | Catalytic rate or formation rate of the product $(Y)$ from $(ES^{\#})$ | $\mu M^{-1}$ | 10.0 | 10.0 | 10.0 | 10.0 | 10.0 |
| $D_h$ | Forward/Backward diffusion rate of the Substrate $(S)$/ Product $(X)$/ Product $(Y)$ | $cm^2min^{-1}$ | 0.0005 | 0.0005 | 0.0005 | 0.0005 | 0.0005 |
| $S_t$ | The total amount of $(S)$ | $\mu M$ | 1000 | 1000 | 1000 | 1000 | 1000 |
| $E_t$ | The total amount of $(E)$ | $\mu M$ | 1000 | 1000 | 1000 | 1000 | 1000 |
| $K$ | Total number of the compartments | - | 50 | 50 | 50 | 50 | 50 |

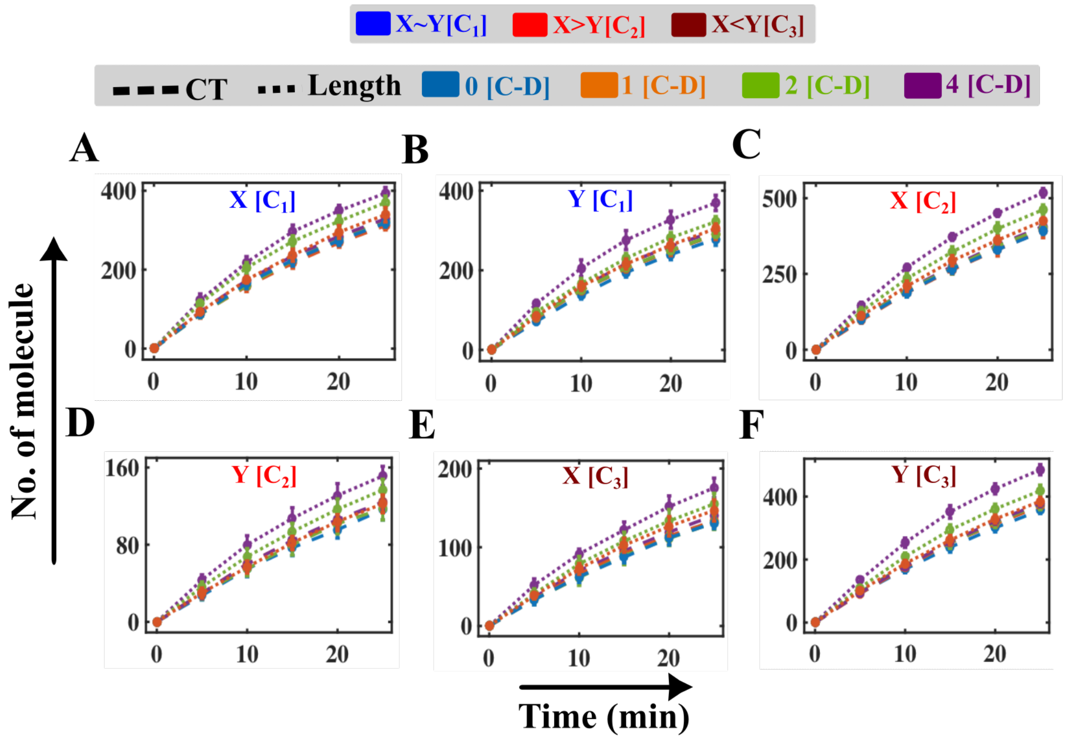

**Fig. S3 Time-course simulation of the product formation altering length (L) for three different scenarios [$C_1$, $C_2$, and $C_3$] for scheme-II.**

Enhancement of the product formation for the increment in the binding rate of the enzyme with the substrate ($k_{bnew} = \frac{k_b}{L}, L = 0.1$) due to the compression of length (L), while performing [C-D] cycles compared to the CT type (when no binding rate $k_b$ has been altered) in case of $C_1$ **(A-B)**, $C_2$ **(C-D),** and $C_3$ **(E-F)**, scheme-II. The rest of the parameters have been kept constant as depicted in Table A5.1.

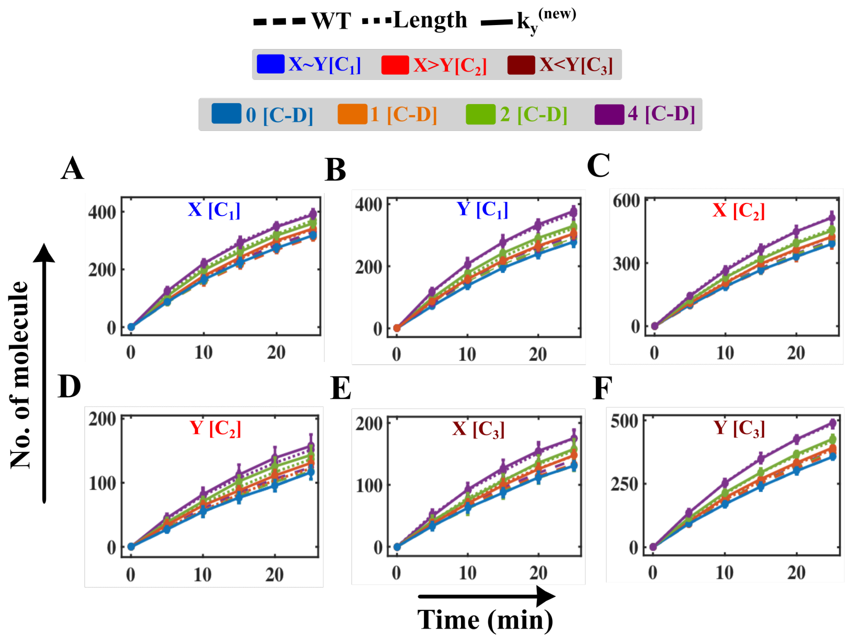

**Fig. S4 Model simulated trajectories of the product altering the catalytic rate $k_y$ for three different scenarios [$C_1$, $C_2$ and $C_3$] for scheme-II.**

The rate of the product formation for the increment in the catalytic rate of the enzyme ($k_y^{new}$[C-D] $= 100 \times k_y$ (CT)), while performing [C-D] cycles compared to the CT type (when no binding rate $k_b$ has been altered) in case of $C_1$ **(A-B)**, $C_2$ **(C-D)** and $C_3$ **(E-F)**, scheme-II. The rest of the parameters have been kept constant as depicted in Table A5.1.

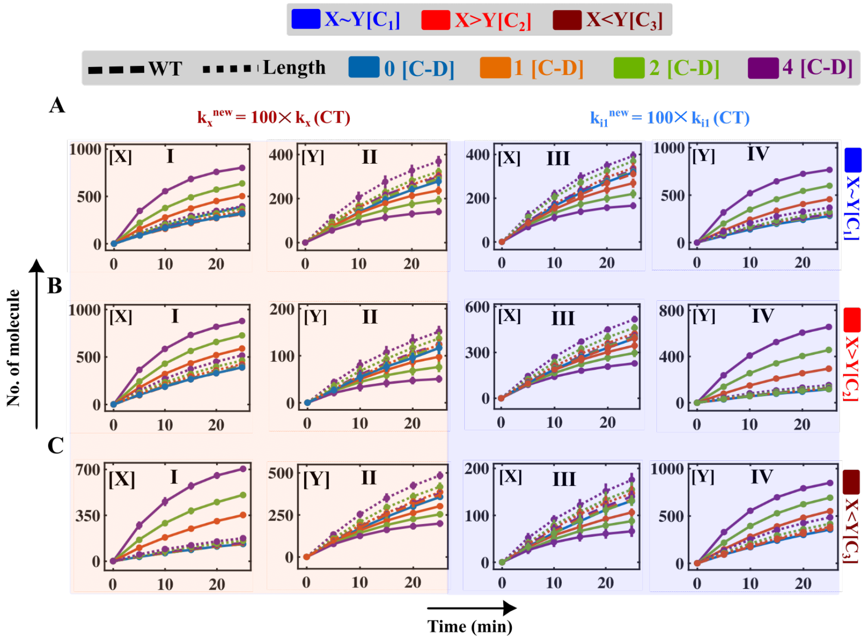

**Fig. S5 Simulated trajectories of the products altering catalytic rate ($k_x$, $k_{i1}$) along with length (L) for three different scenarios [$C_1$, $C_2$ and $C_3$] for scheme-II.**

Comparison of the formation of the product ($X$) and product ($Y$) for CT, changing the catalytic rate along with altering the length. The dashed lines depict the formation of the product ($X$) and product ($Y$), when no catalytic rate has been changed (CT) during performing [C-D] cycles in case of $C_1, C_2$, and $C_3$. The dotted lines represent the enhancement of the product formation for the increment in the binding rate of the enzyme with the substrate ($k_{bnew} = \frac{k_b}{L}, L = 0.1$) due to the compression of the length (L) while performing [C-D] cycles in case of $C_1, C_2$, and $C_3$. The amount of product $X$ (**AI** for case $C_1$, **BI** for case $C_2$, and **CI** for case $C_3$) and product $Y$ (**AII** for $C_1$ case, **BII** for $C_2$ case, and **CII** for $C_3$ case), if the [C-D] cycles accelerate the catalytic rate $k_x$ 100 times compared to CT. The amount of product $X$ (**AIII** for case $C_1$, **BIII** for case $C_2$, and **CIII** for case $C_3$) and product $Y$ (**AIV** for $C_1$ case, **BIV** for $C_2$ case, and **CIV** for $C_3$ case), when the [C-D] cycles fine-tune the conversion rate as ($k_{i1}^{new}$[C-D] =

$100 \times k_{i1}$(CT)).

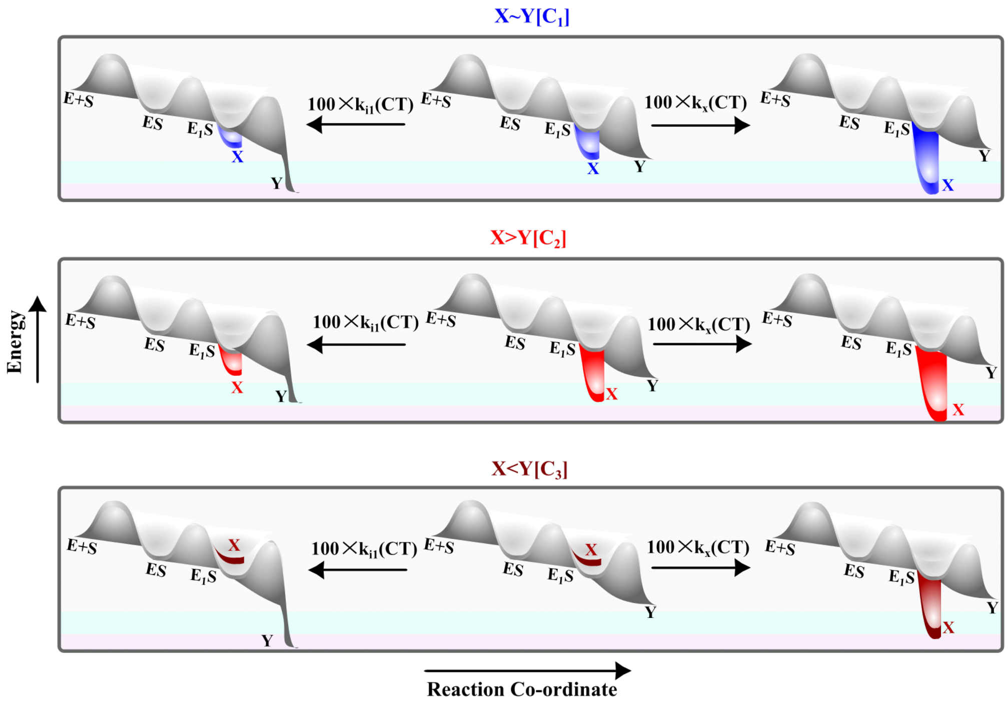

**Fig. S6 Schematic of free energy diagram for the reaction (scheme-II) by changing the reaction fluxes under different conditions.**

The free energy diagram describes the reactions pathway as depicted in the fate of the product X ($k_x$) and Y ($k_{i1}$) formation in Fig. A5.5 under different parametric conditions (Table A5.1).

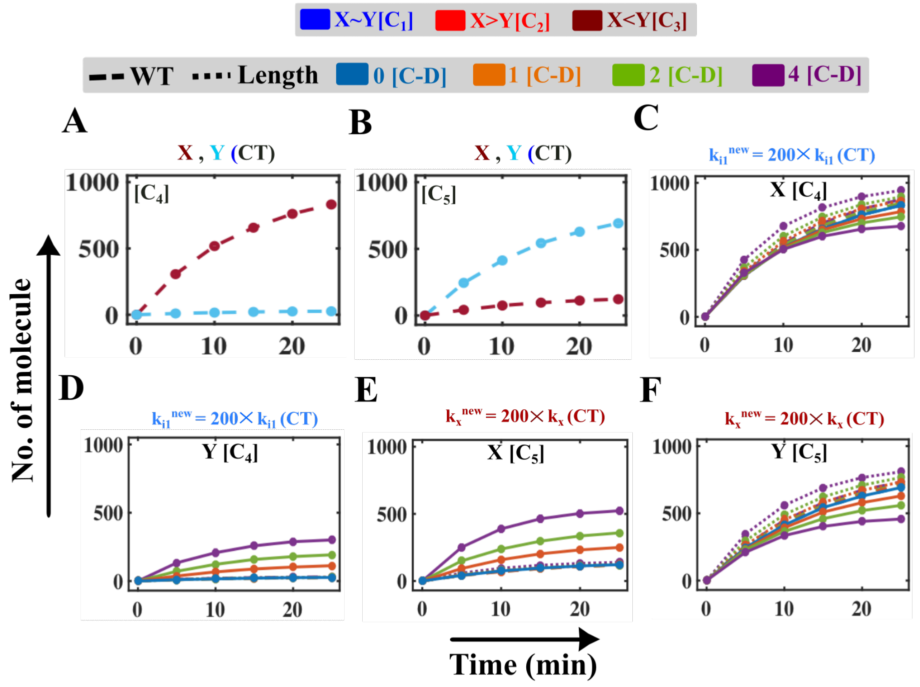

**Fig. S7 Fine-tuning of catalytic rates for case $C_4$ & $C_5$ to alter the formation of products X and Y produced following the mechanism depicted in scheme-II.**

The product formation for scheme-II depicts two extreme scenarios, **A)** where $X >> Y$ ($C_4$) and **B)** $X << Y(C_5)$, without any compression-decompression experiment (CT, 0 [C-D] cycle) using the parameter set provided in Table A5.1. The solid lines in **C)** and **D)** depict the amount of product $X$ and $Y$ in the case of $C_4$, when the [C-D] cycles (1,2 & 4) accelerate the transition rate ($k_{i1}$) almost 200 times. For case $C_5$, the simulated trajectory of $X$ **(E)** and $Y$ **(F)** increasing the catalytic rate ($k_x$) almost 200 folds compare to CT (0 [C-D] cycle). The dashed lines depict the formation of the product ($X$) and product ($Y$), when no catalytic rate has been changed (CT) during performing [C-D] cycle in case of $C_4$ and $C_5$ (Table A5.1). The dotted lines represent the enhanced product formation for the increment in the binding rate of the enzyme with the substrate ($k_{bnew} = \frac{k_b}{L}, L = 0.1$) due to the compression of length (L) while performing [C-D] cycle in case of $C_4$ and $C_5$.

#### Text S6 Observation of the kinetics of the biochemical reaction following mechanisms like scheme-III

**Table S7 Abbreviation of different variables involved in the scheme-III.**

| Symbol | Description (in Conc., $\mu M$) |
|---|---|
| $E$ | The free amount of enzyme |
| $S_1$ | The free amount of substrate $S_1$ |
| $S_2$ | The free amount of another substrate $S_2$ |
| $ES^1$ | The enzyme-substrate complex |
| $ES^{\#\#}$ | Another enzyme-substrate conformational state |
| $Y$ | The amount of product |
| $X$ | The amount of another product |

**Table 8 Description of all the elementary reactions for the proposed scheme-III.**

| Number | Reaction | The propensity of the reaction in $j^{th}$ compartment |
|---|---|---|
| 1 | $E(j) + S_1(j) \rightarrow ES^1(j)$ | $a\_nu(1,j) = k_b \times E(j) \times S_1(j)$ |

| | | |
|---|---|---|
| 2 | $ES^1(j) \rightarrow E(j) + S_1(j)$ | $a\_nu(2,j) = k_b \times ES^1(j)$ |
| 3 | $ES^1(j) + S_2 \rightarrow ES^{\#\#}(j)$ | $a\_nu(3,j) = k_{i1} \times ES^1(j) \times S_2(j)$ |
| 4 | $ES^{\#\#}(j) \rightarrow ES^1(j) + S_2$ | $a\_nu(4,j) = k_{i2} \times ES^{\#\#}(j)$ |
| 5 | $ES^1(j) \rightarrow X(j)$ | $a\_nu(5,j) = k_x \times ES^1(j)$ |
| 6 | $ES^{\#\#}(j) \rightarrow Y(j)$ | $a\_nu(6,j) = k_y \times ES^{\#\#}(j)$ |
| 7 | $S_1(j) \rightarrow S_1(j+1)$ | $a\_nu(7,j) = D_{hfs} \times S_1(j), j = 1\ to\ (K-1)$ |
| 8 | $S_1(j) \rightarrow S_1(j-1)$ | $a\_nu(8,j) = D_{hbs} \times S_1(j), j = 2\ to\ K$ |
| 9 | $S_2(j) \rightarrow S_2(j+1)$ | $a\_nu(7,j) = D_{hfs} \times S_2(j), j = 1\ to\ (K-1)$ |
| 10 | $S_2(j) \rightarrow S_2(j-1)$ | $a\_nu(8,j) = D_{hbs} \times S_2(j), j = 2\ to\ K$ |
| 11 | $X(j) \rightarrow X(j+1)$ | $a\_nu(9,j) = D_{hfp} \times X(j), j = 1\ to\ (K-1)$ |
| 12 | $X(j) \rightarrow X(j-1)$ | $a\_nu(10,j) = D_{hbp} \times X(j), j = 2\ to\ K$ |
| 13 | $Y(j) \rightarrow Y(j+1)$ | $a\_nu(11,j) = D_{hfp} \times Y(j), j = 1\ to\ (K-1)$ |
| 14 | $Y(j) \rightarrow Y(j-1)$ | $a\_nu(12,j) = D_{hbp} \times Y(j), j = 2\ to\ K$ |
| **Algebraic Relation** | | |
| $E_t(j) = E(j) + ES^1(j) + ES^{\#\#}(j)$ | | |
| $S_{1t}(j) = S_1(j) + ES^1(j) + ES^{\#\#}(j) + X(j)$ | | |
| $S_{2t}(j) = S_2(j) + ES^{\#\#}(j) + Y(j)$ | | |

**Table S9 Description of the parameters associated with the scheme - III for different cases.**

| Symbol | Description | Unit | $C_1$ | $C_2$ | $C_3$ | $C_4$ | $C_5$ |
|---|---|---|---|---|---|---|---|
| $k_b$ | Binding rate of the enzyme $(E)$ with the substrate $(S_1)$ | $\mu M^{-1} min^{-1}$ | 0.002 | 0.002 | 0.002 | 0.002 | 0.002 |
| $k_d$ | The dissociation rate of the enzyme-substrate complex $(ES^1)$ | $min^{-1}$ | 10 | 10 | 10 | 10 | 10 |
| $k_{i1}$ | Binding rate of complex $(ES^1)$ with the substrate $(S_2)$ | $\mu M^{-1} min^{-1}$ | 0.5 | 0.5 | 1.2 | 0.5 | 2 |

| | | | | | | | |
|---|---|---|---|---|---|---|---|
| $k_{i2}$ | The conversion rate of complex $(ES^{\#\#})$ to $(ES^1)$ | $\mu M^{-1}$ | 1.0 | 1.0 | 1.0 | 1.0 | 1.0 |
| $k_x$ | Catalytic rate or the formation rate of the product $(X)$ from $(ES^1)$ | $\mu M^{-1}$ | 4.0 | 12.0 | 4.0 | 100.0 | 0.4 |
| $k_y$ | Catalytic rate or formation rate of the product $(Y)$ from $(ES^{\#\#})$ | $\mu M^{-1}$ | 1.0 | 1.0 | 1.0 | 1.0 | 1.0 |
| $D_h$ | Forward/ Backward diffusion rate of the Substrate $(S_1)$/Substrate $(S_2)$ Product $(X)$/ Product $(Y)$ | $cm^2 min^{-1}$ | 0.0005 | 0.0005 | 0.0005 | 0.0005 | 0.0005 |
| $S_{1t}$ | The total amount of $(S_1)$ | $\mu M$ | 0.0005 | 0.0005 | 1000 | 1000 | 1000 |
| $S_{2t}$ | The total amount of $(S_2)$ | $\mu M$ | 0.0005 | 0.0005 | 1000 | 1000 | 1000 |
| $E_t$ | The total amount of $(E)$ | $\mu M$ | 0.0005 | 0.0005 | 1000 | 1000 | 1000 |
| $K$ | Total number of compartment | − | 50 | 50 | 50 | 50 | 50 |

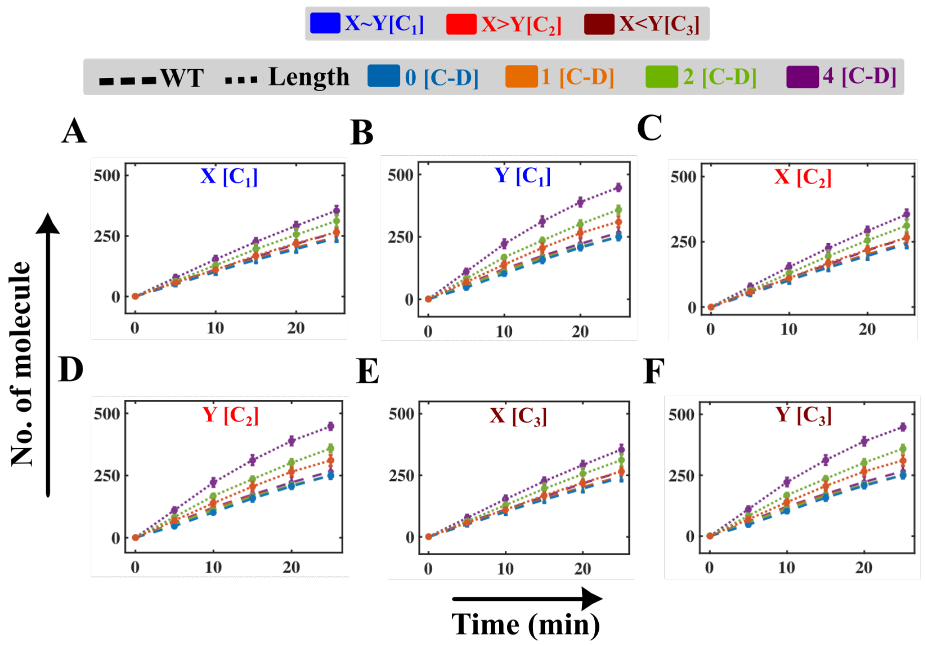

**Fig. S8 Time-course simulation of the product formation altering length (L) for three different scenarios [$C_1$, $C_2$ and $C_3$] for scheme-III.**

Enhancement of the product formation for the increment in the binding rate of the enzyme with the substrate ($k_{bnew} = \frac{k_b}{L}, L = 0.1$) due to the compression of length (L), while performing [C-D] cycles compared to the CT type (when no binding rate $k_b$ has been altered) in case of $C_1$ **(A-B)**, $C_2$ **(C-D)** and $C_3$ **(E-F)**, scheme-III. The rest of the parameters have been kept constant as depicted in Table A5.2.

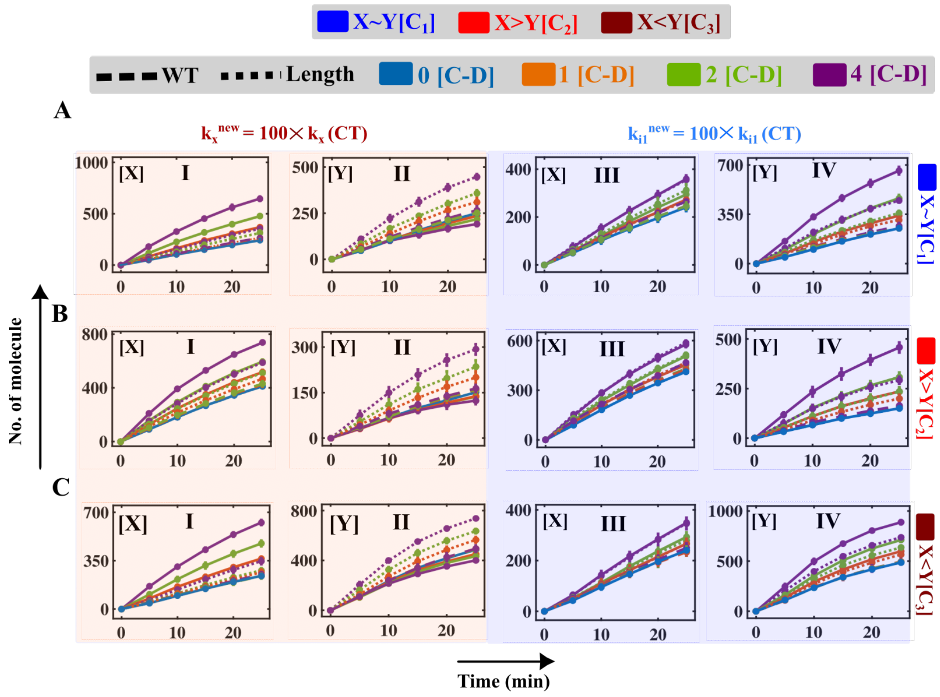

**Fig. S9 Simulated trajectories of the products altering catalytic rate ($k_x, k_{i1}$) along with length (L) for three different scenarios [$C_1$, $C_2$ and $C_3$] for schematic-III.**

Comparison of the formation of the product ($X$) and product ($Y$) for CT, changing the catalytic rate along with altering the length. The dashed lines depict the formation of the product ($X$) and product ($Y$), when no catalytic rate has been changed (CT) during performing [C-D] cycles in case of $C_1, C_2$ and $C_3$. The dotted lines represent the enhancement of the product formation for the increment in the binding rate of the enzyme with the substrate ($k_{bnew} = \frac{k_b}{L}, L = 0.1$) due to the compression of the length (L) while performing [C-D] cycles in case of $C_1, C_2$ and $C_3$. The amount of product $X$ (**AI** for case $C_1$, **BI** for case $C_2$, and **CI** for case $C_3$) and product $Y$ (**AII** for $C_1$ case, **BII** for $C_2$ case, and **CII** for $C_3$ case), if the [C-D] cycles accelerate the catalytic rate $k_x$ 100 times compared to CT. The amount of product $X$ (**AIII** for case $C_1$, **BIII** for case $C_2$, and **CIII** for case $C_3$) and product $Y$ (**AIV** for $C_1$ case, **BIV** for $C_2$ case, and **CIV** for $C_3$ case), when the [C-D] cycles fine-tune the conversion rate as ($k_{i1}^{new}$[C-D] $= 100 \times k_{i1}$(CT)).

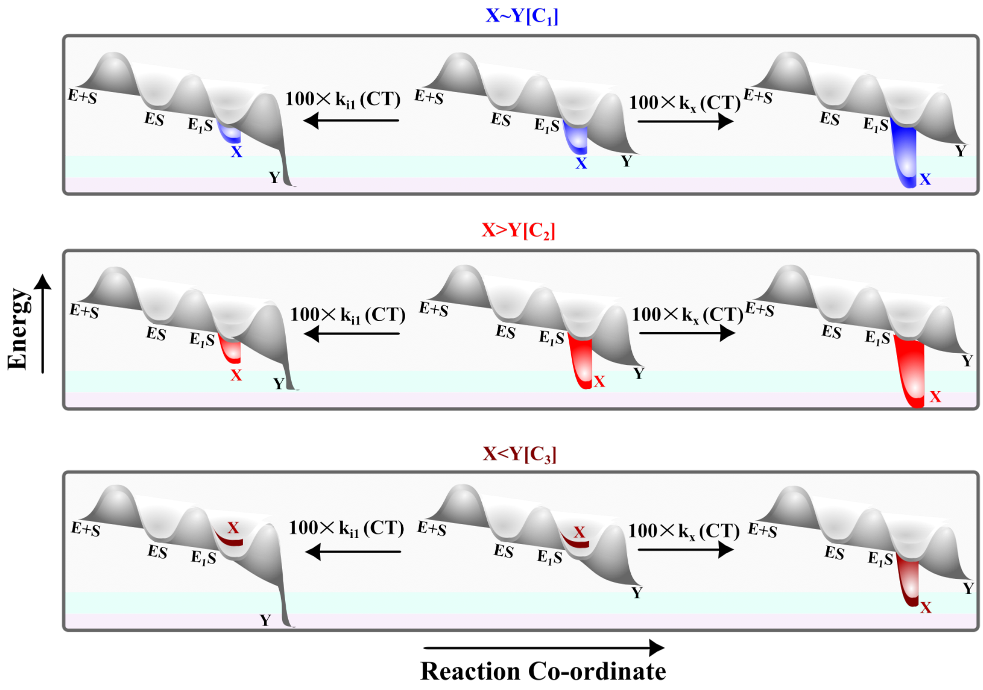

**Fig. S10 Schematic of free energy diagram for the reaction (scheme-III) by changing the reaction fluxes under different conditions.**

The free energy diagram describes the reactions pathway as depicted in the fate of the product X ($k_x$) and Y ($k_{i1}$) formation in Fig. A5.9 under different parametric conditions (Table A5.2).

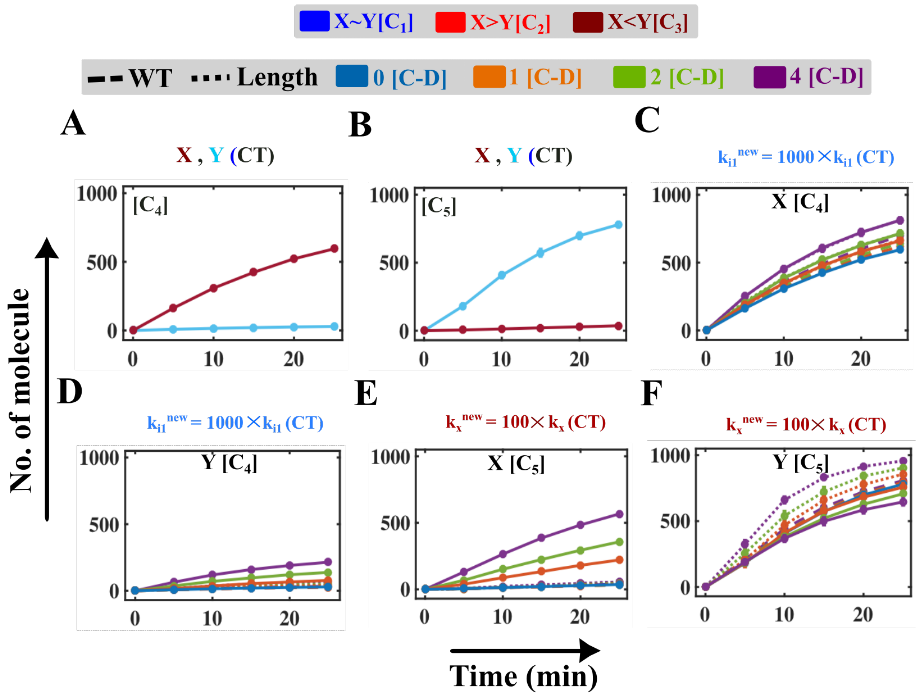

**Fig. S11 Fine-tuning of catalytic processes for case $C_4$ & $C_5$ to alter the formation of products $X$ and $Y$ produced following the mechanism depicted in scheme-III.**

The product formation for scheme-II depicts two extreme scenarios, **A)** where $X >> Y$ ($C_4$) and **B)** $X << Y(C_5)$, without any compression-decompression experiment (CT, 0 [C-D] cycle) using the parameter set provided in Table A5.1. The solid lines in **C)** and **D)** depict the amount of product $X$ and $Y$ in the case of $C_4$**,** when the [C-D] cycles (1,2 & 4) accelerate the transition rate almost ($k_{i1}$) 100 times. For case $C_5$, the simulated trajectory of $X$ **(E)** and $Y$ **(F)** increasing the catalytic rate ($k_x$) almost 300 folds compare to CT (0 [C-D] cycle). The dashed lines depict the formation of the product ($X$) and product ($Y$), when no catalytic rate has been changed (CT) during performing [C-D] cycle in case of $C_4$ and $C_5$ (Table A5.2). The dotted lines represent the enhanced product formation for the increment in the binding rate of the enzyme with the substrate ($k_{bnew} = \frac{k_b}{L}, L = 0.1$) due to the compression of length (L) while performing [C-D] cycle in case of $C_4$ and $C_5$.

#### Text S7 Observation of the kinetics of the biochemical reaction following mechanisms like scheme-IV

**Table S10 Description of all the elementary reactions for the proposed scheme-IV.**

| **Number** | **Symbol** | **Description (in Conc., $\mu mol$)** |
|---|---|---|
| 1 | $E$ | The free amount of enzyme |
| 2 | $S_1$ | The free amount of substrate |
| 3 | $S_2$ | The free amount of another substrate |
| 4 | $ES^1$ | The enzyme-substrate complex |
| 5 | $ES^2$ | Another enzyme-substrate complex |
| 6 | $ES^{\#\#}$ | Another hetero-trimeric enzyme-substrate complex |
| 7 | $Y$ | The amount of product |
| 8 | $X$ | The amount of another product |
| **Number** | **Reaction** | **The propensity of the reaction in $j^{th}$ compartment** |
| 1 | $E(j) + S_1(j) \rightarrow ES^1(j)$ | $a\_nu(1, j) = k_{b1} \times E(j) \times S_1(j)$ |
| 2 | $ES^1(j) \rightarrow E(j) + S_1(j)$ | $a\_nu(2, j) = k_{d1} \times ES^1(j)$ |
| 3 | $E(j) + S_2(j) \rightarrow ES^2(j)$ | $a\_nu(3, j) = k_{b2} \times E(j) \times S_2(j)$ |
| 4 | $ES^2(j) \rightarrow E(j) + S_2(j)$ | $a\_nu(4, j) = k_{d2} \times ES^2(j)$ |
| 5 | $ES^1(j) + S_2 \rightarrow ES^{\#\#}(j)$ | $a\_nu(5, j) = k_{i1} \times ES^1(j) \times S_2(j)$ |
| 6 | $ES^{\#\#}(j) \rightarrow ES_1(j) + S_2$ | $a\_nu(6, j) = k_{i2} \times ES^{\#\#}(j)$ |
| 7 | $ES^2(j) + S_1 \rightarrow ES^{\#\#}(j)$ | $a\_nu(7, j) = k_{ii1} \times ES^2(j) \times S_1(j)$ |
| 8 | $ES^{\#\#}(j) \rightarrow ES^2(j) + S_1$ | $a\_nu(8, j) = k_{ii2} \times ES^{\#\#}(j)$ |
| 9 | $ES^1(j) \rightarrow X(j) + E(j)$ | $a\_nu(9, j) = k_x \times ES^1(j)$ |
| 10 | $ES^{\#\#}(j) \rightarrow Y(j) + ES^1(j)$ | $a\_nu(10, j) = k_y \times ES^{\#\#}(j)$ |
| 11 | $S_1(j) \rightarrow S_1(j+1)$ | $a\_nu(11, j) = D_{hfs} \times S_1(j), j = 1\ to\ (K-1)$ |
| 12 | $S_1(j) \rightarrow S_1(j-1)$ | $a\_nu(12, j) = D_{hbs} \times S_1(j), j = 2\ to\ K$ |
| 13 | $S_2(j) \rightarrow S_2(j+1)$ | $a\_nu(13, j) = D_{hfs} \times S_2(j), j = 1\ to\ (K-1)$ |
| 14 | $S_2(j) \rightarrow S_2(j-1)$ | $a\_nu(14, j) = D_{hbs} \times S_2(j), j = 2\ to\ K$ |
| 15 | $X(j) \rightarrow X(j+1)$ | $a\_nu(15, j) = D_{hfp} \times X(j), j = 1\ to\ (K-1)$ |
| 16 | $X(j) \rightarrow X(j-1)$ | $a\_nu(16, j) = D_{hbp} \times X(j), j = 2\ to\ K$ |
| 17 | $Y(j) \rightarrow Y(j+1)$ | $a\_nu(17, j) = D_{hfp} \times Y(j), j = 1\ to\ (K-1)$ |
| 18 | $Y(j) \rightarrow Y(j-1)$ | $a\_nu(18, j) = D_{hbp} \times Y(j), j = 2\ to\ K$ |
| **Algebraic Relation** | | |
| $E_t(j) = E(j) + ES^1(j) + ES^2(j) + ES^{\#\#}(j)$ | | |

| |
|---|
| $S_{1t}(j) = S_1(j) + ES^1(j) + ES^{\#\#}(j) + X(j)$ |
| $S_{2t}(j) = S_2(j) + ES^2(j) + ES^{\#\#}(j) + Y(j)$ |

**Table S11 Description of the parameters associated with the scheme-IV for different cases**

| Symbol | Description | Unit | [$T_1$] | [$T_2$] | [$T_3$] | [$T_4$] | [$T_5$] |
|---|---|---|---|---|---|---|---|
| $k_{b1}$ | Binding rate of the enzyme ($E$) with the substrate ($S_1$) | $\mu M^{-1} min^{-}$ | 0.001 | 0.002 | 0.002 | 0.002 | 0.002 |
| $k_{d1}$ | The dissociation rate of the enzyme-substrate complex ($ES^1$) | $min^{-1}$ | 10 | 10 | 10 | 10 | 10 |
| $k_{b2}$ | Binding rate of the enzyme ($E$) with the substrate ($S_2$) | $\mu M^{-1} min^{-}$ | 0.0015 | 0.002 | 0.002 | 0.002 | 0.002 |
| $k_{d2}$ | The dissociation rate of the enzyme-substrate complex ($ES^2$) | $\mu M^{-1}$ | 1 | 10 | 10 | 10 | 10 |
| $k_{i1}$ | Binding rate of complex ($ES^1$) with the substrate ($S_2$) | $\mu M^{-1} min^{-}$ | 0.2 | 0.5 | 1.2 | 0.5 | 2 |
| $k_{i2}$ | The dissociation rate of complex ($ES^{\#\#}$) | $\mu M^{-1}$ | 1.0 | 1.0 | 1.0 | 1.0 | 1.0 |
| $k_{ii1}$ | Binding rate of complex ($ES^2$) with the substrate ($S_1$) | $\mu M^{-1} min^{-}$ | 0.05 | 0.5 | 1.2 | 0.5 | 2 |
| $k_{ii2}$ | The dissociation rate of complex ($ES^{\#\#}$) | $\mu M^{-1}$ | 0.1 | 1.0 | 1.0 | 1.0 | 1.0 |
| $k_x$ | Catalytic rate or formation rate of the product ($X$) from ($ES^1$) | $\mu M^{-1}$ | 3.0 | 12.0 | 4.0 | 100.0 | 0.4 |

| | | | | | | | |
|---|---|---|---|---|---|---|---|
| $k_y$ | Catalytic rate or formation rate of the product $(Y)$ from $(ES^{\#\#})$ | $\mu M^{-1}$ | 1.0 | 1.0 | 1.0 | 1.0 | 1.0 |
| $D_h$ | Forward/ Backward diffusion rate of the substrate $(S_1)$/ substrate $(S_2)$ product $(X)$/ product $(Y)$ | $cm^2min^{-1}$ | 0.000 5 | 0.000 5 | 0.000 5 | 0.000 5 | 0.000 5 |
| $S_{1t}$ | The total amount of $(S_1)$ | $\mu M$ | | | 1000 | 1000 | 1000 |
| $S_{2t}$ | The total amount of $(S_2)$ | $\mu M$ | | | 1000 | 1000 | 1000 |
| $E_t$ | The total amount of $(E)$ | $\mu M$ | | | 1000 | 1000 | 1000 |
| $K$ | Total number of compartments | - | | | 50 | 50 | 50 |

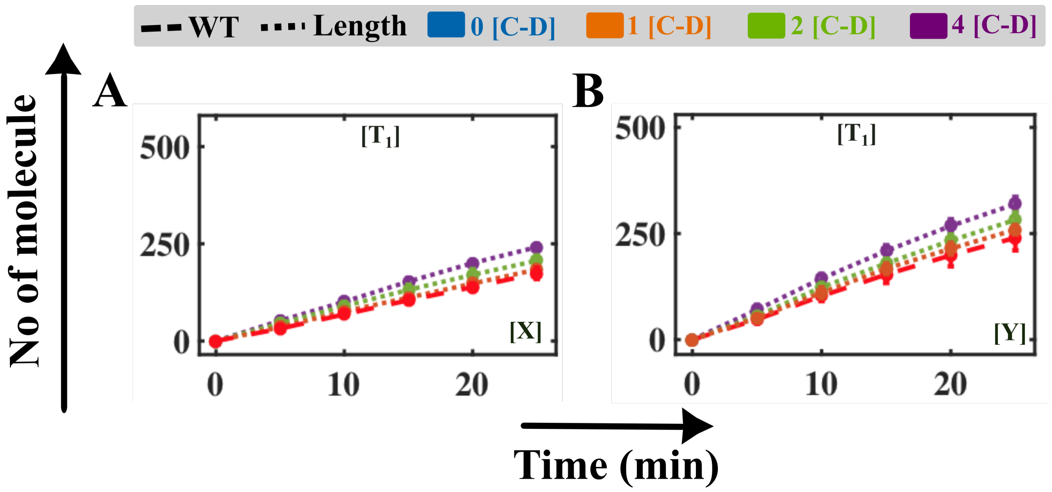

**Fig. S12 Kinetics of the evolution of the product in case of both the substrate has comparable binding rate ($T_1$) for scheme-IV.**

Enhancement of the product $X$ **(A)** and $Y$ **(B)** formation for the increment in the binding rate of the enzyme with the substrate $\left[k_{b1}(new) = \left({}^{k_{b1}}/_{L}\right), k_{b2}(new) = \left({}^{k_{b2}}/_{L}\right), L = 0.1\right]$ due to the compression of length (L) while performing the [C-D] cycle compared to the CT type (when no binding rate $K_b$ has been altered in scheme-IV, dashed lines).

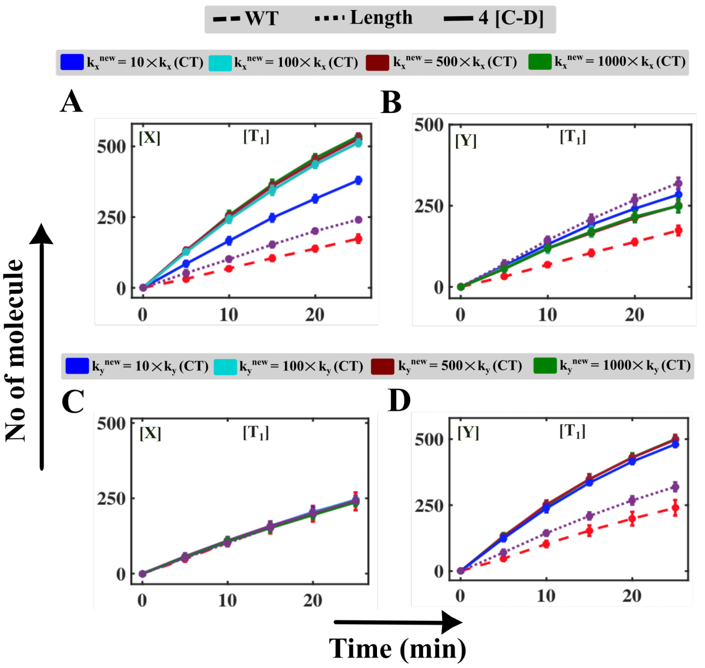

**Fig. S13 Evolution of the products altering the catalytic rates to different extents in case of ( $T_1$) following scheme-IV.**

**A, B)** Solid lines represent the simulated trajectories of the products $X$ and $Y$, enhancing the catalytic rate $k_x$ during the [C-D] cycles compared to the CT type (when no binding rate $K_b$ has been altered scheme-IV). **C, D)** Solid lines represent the simulated time profiles of product $X$ and $Y$ increasing catalytic rate of Y $k_y(new) = 10 \times k_y; 50 \times k_y; 100 \times k_y, 1000 \times k_y$]. Dotted lines represent the product $X$ and $Y$ formation for the length (L) effect while performing the [C-D] cycle compared to the CT type (when no binding rate $k_b$ has been altered in scheme-IV, dashed lines)).

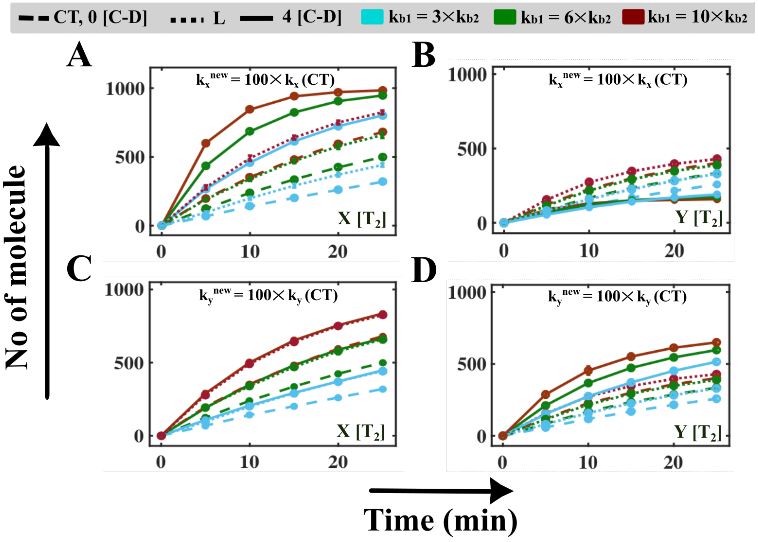

**Fig. S14 Kinetics of the evolution of product altering catalytic $\left(k_x, k_y\right)$ for $(T_1)$ in the case of scheme-IV.**

**A)** Enhancement of the catalytic rate $(k_x(new) = 100 \times k_x(CT))$ during 4 [C-D] cycle results in the increment of the product $X$ and **B)** reduction of the product $Y$ significantly. **C)** Simulated time profiles by altering the catalytic rate $\left(k_y(new) = 100 \times k_y(CT)\right)$ during 4 [C-D] cycles, which indicates the negligible reduction of $X$ compared to the length effect and **D)** moderate enhancement of the product $Y$. The simulation has been performed by increasing the binding rate as $[k_{b1}(new) = 3 \times k_{b2}; 6 \times k_{b2}; 10 \times k_{b2}]$ keeping all other rates constant same (Table A5.3) and considering the length effect as for $T_2$. The dotted lines represent the model predicted profile at 0 [C-D] cycle (CT, when no other rates have been altered).

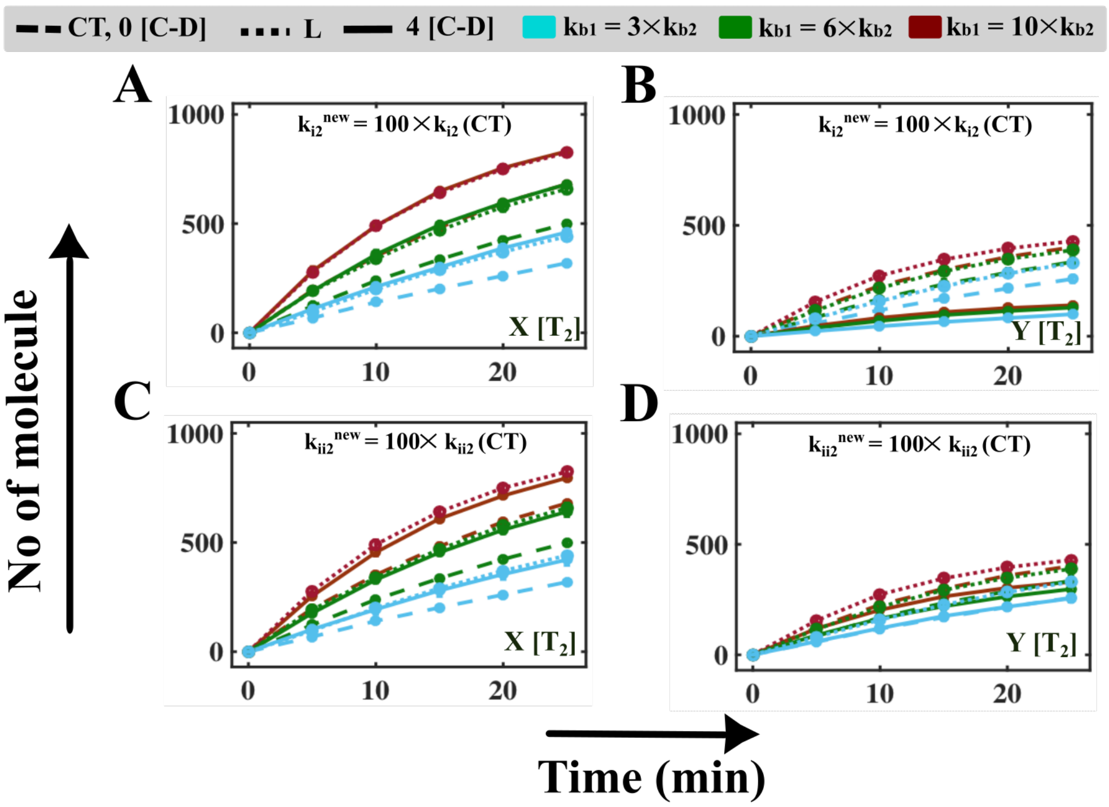

**Fig. S15 Kinetics of the evolution of product altering conversion rate $(k_{i2}, k_{ii2})$ for $T_2$ in the case of scheme-IV.**

**A)** Enhancement of the conversion rate $(k_{i2}(new) = 100 \times k_{i2}(CT))$ during 4 [C-D] cycle results in the increment of the product $X$ and **B)** reduction of the product $Y$ significantly. **C)** Simulated time profiles by altering the catalytic rate $(k_{ii2}(new) = 100 \times k_{ii2}(CT))$ during 4 [C-D] cycles, which indicates the negligible reduction of $X$ compared to the length effect and **D)** moderate enhancement of the product $Y$. The simulation has been performed by increasing the binding rate as $[k_{b1}(new) = 3 \times k_{b2}; 6 \times k_{b2}; 10 \times k_{b2}]$ keeping all other rates constant same (Table A5.3) and considering the length effect as for $T_2$. The dotted lines represent the model predicted profile at 0 [C-D] cycle (CT, when no other rates have been altered), and the solid lines indicate the trajectory at 4 [C-D] cycle.

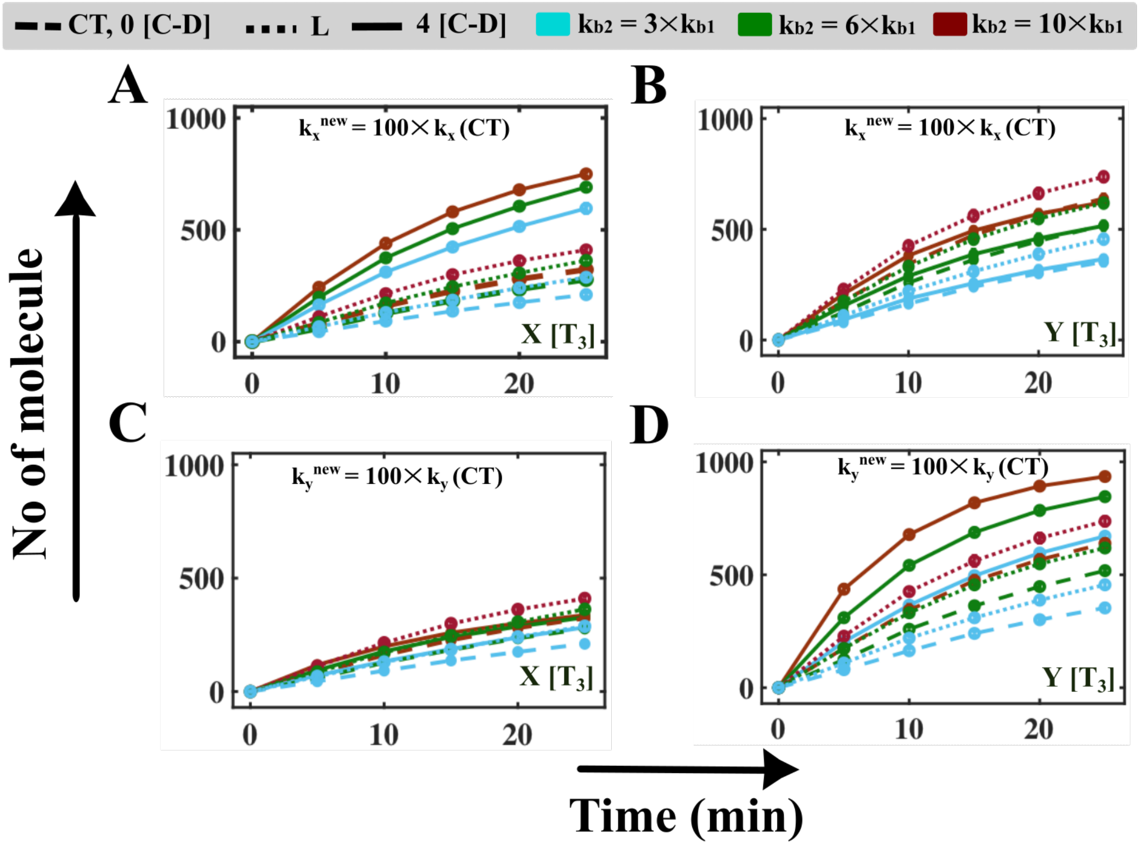

**Fig. S16 Kinetics of the evolution of product altering catalytic $(k_x, k_y)$ for (T3) in the case of scheme-IV.**

**A)** Enhancement of the catalytic rate $(k_x(new) = 100 \times k_x(CT))$ during 4 [C-D] cycle results in the increment of the product $X$ and **B)** reduction of the product $Y$ significantly. **C)** Simulated time profiles by altering the catalytic rate $(k_y(new) = 100 \times k_y(CT))$ during 4 [C-D] cycles, which indicates the negligible reduction of $X$ compared to the length effect and **D)** moderate enhancement of the product $Y$. The simulation has been performed by increasing the binding rate as $[k_{b2}(new) = 3 \times k_{b1}; 6 \times k_{b1}; 10 \times k_{b1}]$ keeping all other rates constant same (Table A5.3) and considering the length effect as for $T_3$. The dotted lines represent the model predicted profile at 0 [C-D] cycle (CT, when no other rates have been altered), and the solid lines indicate the trajectory at 4 [C-D] cycle.

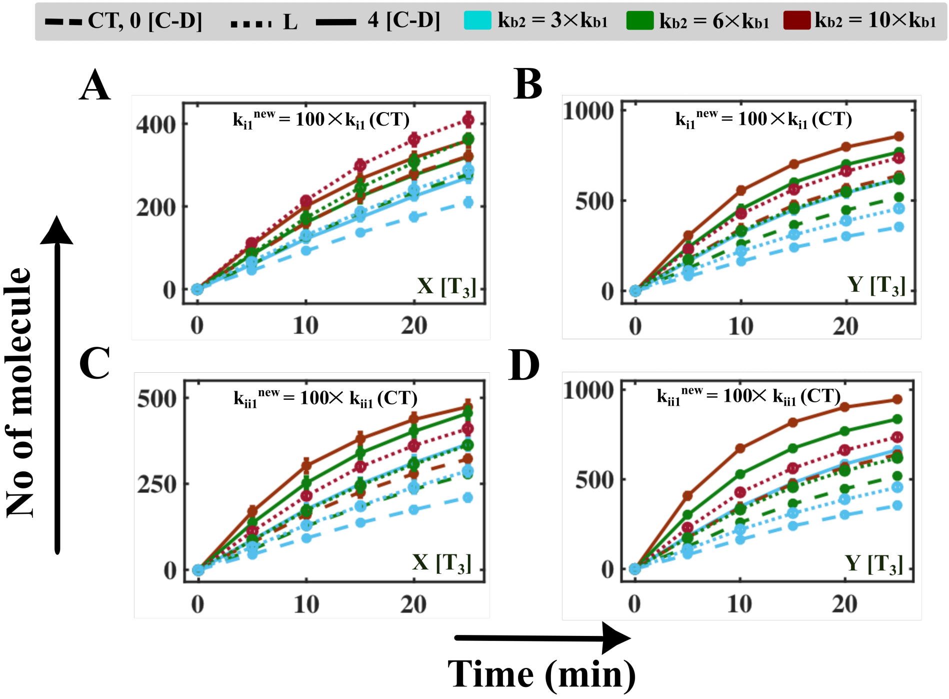

**Fig. S17 Simulated trajectories of the product formation altering conversion rate ($k_{i1}, k_{ii1}$) for $T_3$ in the case of scheme-IV.**

**A)** Enhancement of the catalytic rate ($k_{i1}(new) = 100 \times k_{i1}(CT)$) during 4 [C-D] cycle results in the increment of the product $X$ and **B)** reduction of the product $Y$ significantly. **C)** Simulated time profiles by altering the catalytic rate ($k_{ii1}(new) = 100 \times k_{ii1}(CT)$) during 4 [C-D] cycles, which indicates the negligible reduction of $X$ compared to the length effect and **D)** moderate enhancement of the product $Y$. The simulation has been performed by increasing the binding rate as [$k_{b2}(new) = 3 \times k_{b1}; 6 \times k_{b1}; 10 \times k_{b1}$] keeping all other rates constants similar (Table A5.3) and considering the length effect as for $T_3$. The dotted lines represent the model predicted profile at 0 [C-D] cycle (CT, when no other rates have been altered), and the solid lines indicate the trajectories at 4 [C-D] cycle.

#### Text S8 Observation of the kinetics of the biochemical reaction following mechanisms like scheme-V

**Table S12 Description of all the variables and elementary reactions for the proposed scheme-V**

| Number | Symbol | Description (in Conc., $\mu mol$) |
|---|---|---|
| 1 | $S$ | The amount of free substrate |

| | | |
|---|---|---|
| 2 | $E_1$ | The amount of free enzyme |
| 3 | $E_2$ | The free amount of another enzyme |
| 4 | $E^1S$ | The enzyme-substrate complex |
| 5 | $E^2S$ | Another enzyme-substrate complex |
| 6 | $E^{\#1}S$ | Another enzyme-substrate complex originates from $E^1S$ |
| 7 | $E^{\#2}S$ | Another enzyme-substrate complex originates from $E^2S$ |
| 8 | $X$ | The amount of product |
| 9 | $Y$ | The amount of another product |
| **Number** | **Reaction** | **The propensity of the reaction in $j^{th}$ compartment** |
| 1 | $E_1(j) + S(j) \rightarrow E^1S(j)$ | $a\_nu(1,j) = k_{b1} \times E_1(j) \times S(j)$ |
| 2 | $E^1S(j) \rightarrow E_1(j) + S(j)$ | $a\_nu(2,j) = k_{d1} \times E^1S(j)$ |
| 3 | $E_2(j) + S(j) \rightarrow E^2S(j)$ | $a\_nu(3,j) = k_{b2} \times E_2(j) \times S(j)$ |
| 4 | $E^2S(j) \rightarrow E_2(j) + S(j)$ | $a\_nu(4,j) = k_{d2} \times E^2S(j)$ |
| 5 | $E^1S(j) \rightarrow E^{\#1}S(j)$ | $a\_nu(5,j) = k_{i1} \times E^1S(j)$ |
| 6 | $E^{\#1}S(j) \rightarrow E^1S(j)$ | $a\_nu(6,j) = k_{i2} \times E^{\#1}S(j)$ |
| 7 | $E^2S(j) \rightarrow E^{\#2}S(j)$ | $a\_nu(7,j) = k_{ii1} \times E^2S(j)$ |
| 8 | $E^{\#2}S(j) \rightarrow E^2S(j)$ | $a\_nu(8,j) = k_{ii2} \times E^{\#2}S(j)$ |
| 9 | $E^{\#1}S(j) \rightarrow X(j) + E(j)$ | $a\_nu(9,j) = k_x \times E^{\#1}S(j)$ |
| 10 | $E^{\#2}S(j) \rightarrow Y(j) + E(j)$ | $a\_nu(10,j) = k_y \times E^{\#2}S(j)$ |
| 11 | $S(j) \rightarrow S(j+1)$ | $a\_nu(11,j) = D_{hfs} \times S(j), j = 1\ to\ (K-1)$ |
| 12 | $S(j) \rightarrow S(j-1)$ | $a\_nu(12,j) = D_{hbs} \times S(j), j = 2\ to\ K$ |
| 13 | $X(j) \rightarrow X(j+1)$ | $a\_nu(13,j) = D_{hfp} \times X(j), j = 1\ to\ (K-1)$ |
| 14 | $X(j) \rightarrow X(j-1)$ | $a\_nu(14,j) = D_{hbp} \times X(j), j = 2\ to\ K$ |
| 15 | $Y(j) \rightarrow Y(j+1)$ | $a\_nu(15,j) = D_{hfp} \times Y(j), j = 1\ to\ (K-1)$ |
| 16 | $Y(j) \rightarrow Y(j-1)$ | $a\_nu(16,j) = D_{hbp} \times Y(j), j = 2\ to\ K$ |
| **Algebraic Relation** | | |
| $E_{1t}(j) = E_1(j) + E^1S(j) + E^{\#1}S(j)$ | | |

| |
|---|
| $E_{2t}(j) = E_2(j) + E^2S(j) + E^{\#2}S(j)$ |
| $S_t(j) = S(j) + E^1S(j) + E^{\#1}S(j) + E^2S(j) + E^{\#2}S(j) + X(j) + Y(j)$ |

**Table S13 Description of the parameters associated with the scheme-V for different cases.**

| Symbol | Description | Unit | [T₁] | [T₂] | [T₃] |
|---|---|---|---|---|---|
| $k_{b1}$ | Binding rate of the enzyme $E_1$ with the substrate $S$ | $\mu M^{-1}min^{-1}$ | 0.001 | 0.002/0.005 | 0.001 |
| $k_{d1}$ | The dissociation rate of the enzyme-substrate complex $E^1S$ | $min^{-1}$ | 10 | 10 | 10 |
| $k_{b2}$ | Binding rate of the enzyme $E_2$ with the substrate $S$ | $\mu M^{-1}min^{-1}$ | 0.001 | 0.001 | 0.002/0.005 |
| $k_{d2}$ | The dissociation rate of the enzyme-substrate complex $E^2S$ | $\mu M^{-1}$ | 10 | 10 | 10 |
| $k_{i1}$ | The conversion rate of complex $E^1S$ to complex $E^{\#1}S$ | $\mu M^{-1}$ | 2 | 2 | 2 |
| $k_{i2}$ | The conversion rate of complex $E_1S_1$ to complex $E^1S$ | $\mu M^{-1}$ | 0.1 | 0.1 | 0.1 |
| $k_{ii1}$ | The conversion rate of complex $E^2S$ to complex $E^{\#2}S$ | $\mu M^{-1}$ | 2 | 2 | 2 |
| $k_{ii2}$ | The conversion rate of complex $E^{\#2}S$ to complex $E^2S$ | $\mu M^{-1}$ | 0.1 | 0.1 | 0.1 |
| $k_x$ | Catalytic rate or | $\mu M^{-1}$ | 1.0 | 1.0 | 1.0 |

| | | | | | |
|---|---|---|---|---|---|
| | formation rate of the product $X$ from $E^{\#1}S$ | | | | |
| $k_y$ | Catalytic rate or formation rate of the product $Y$ from $E^{\#2}S$ | $\mu M^{-1}$ | 2.0 | 2.0 | 2.0 |
| $D_h$ | Forward/ Backward diffusion rate of the substrate $S$/ product $X$/ product $Y$ | $cm^2 min^{-1}$ | 0.000 5 | 0.0005 | 0.001 |
| $S_t$ | The total amount of $S$ | $\mu M$ | 2000 | 2000 | 2000 |
| $E_{1t}$ | The total amount of $E_1$ | $\mu M$ | 1000 | 1000 | 1000 |
| $E_{2t}$ | The total amount of $E_2$ | $\mu M$ | 1000 | 1000 | 1000 |
| $K$ | Total number of compartment | - | 50 | 50 | 50 |

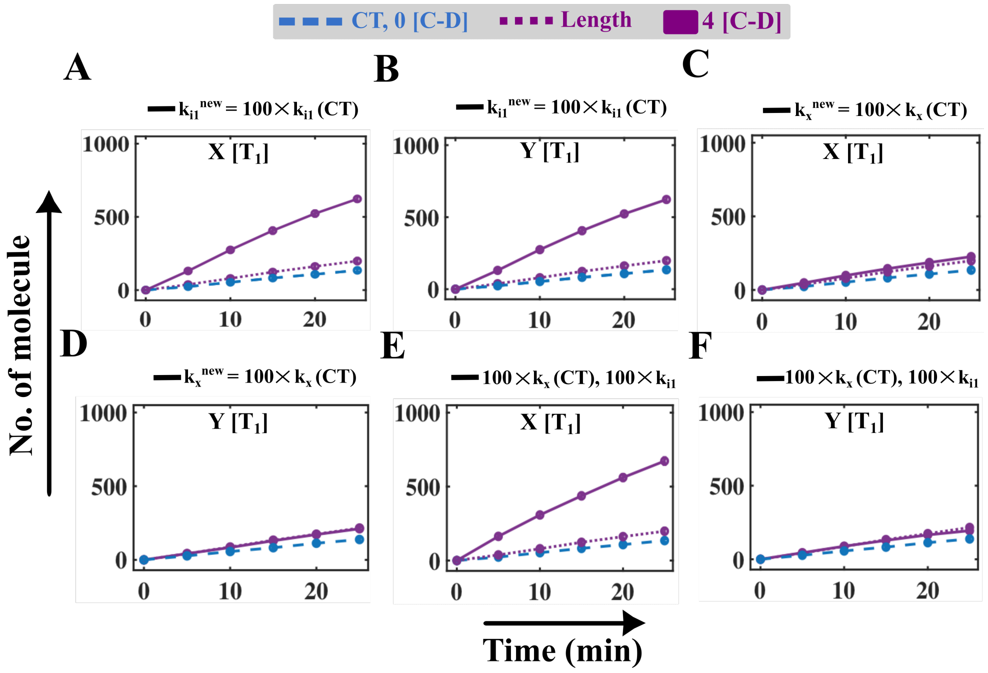

**Fig. S18 Simulated trajectories of the product obtained, when [C-D] cycles altered the catalytic ($k_x$) and conversion rate ($k_{i1}$) in case of scheme-V.**

For case $T_1$, the simulated time profile of product ($X$) and product ($Y$), when [C-D] events (4

[C-D]) fine-tune the intermediate rate $k_{i1}$ and the catalytic rate $k_x$ either individually (solid lines in **A, B, C, and D**). Time evolution of **E)** product ($X$) and **F)** product ($Y$), when [C-D] event (4 [C-D]) alters the intermediate rate $k_{i1}$ and the catalytic rate $k_x$ collectively (solid lines). The simulation has been performed by taking the binding rates similar and keeping all other rate constants the same (Table A5.5). The dashed lines represent the model-predicted profile for CT, and the dotted lines indicate the model-predicted profile altering the length ($k_{b1}/L$, $k_{b2}/L$, $L = 0.1$) at different [C-D] cycles.

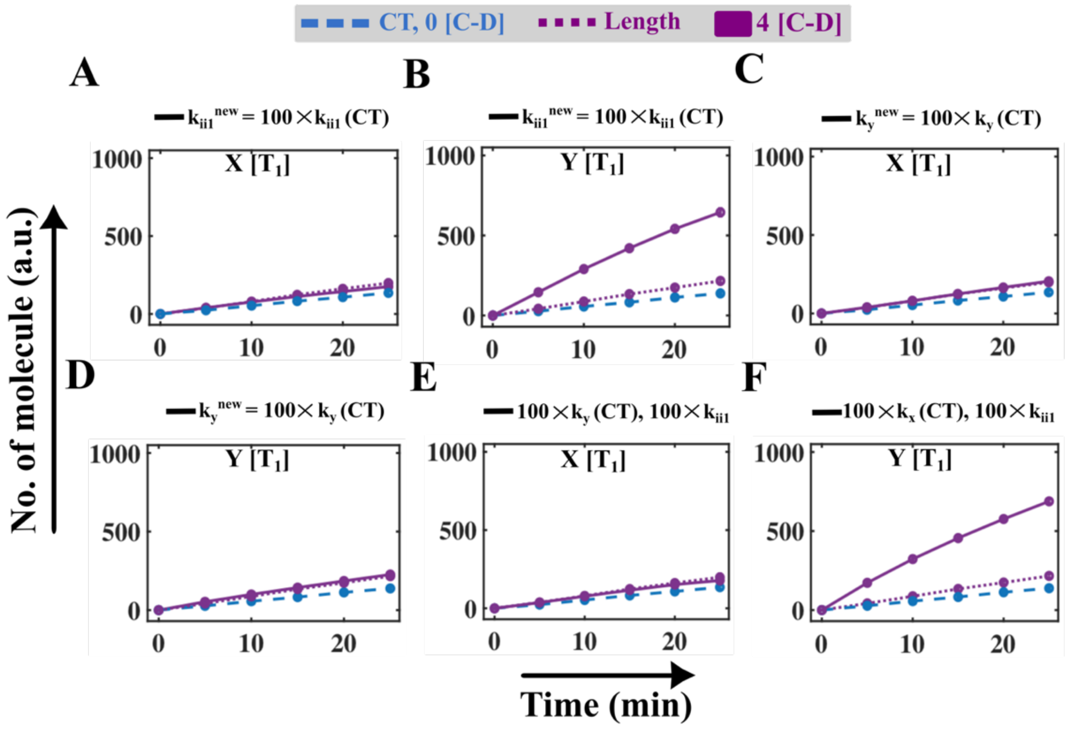

**Fig. A5.19 Simulated trajectories of the product obtained, when [C-D] cycles altered the catalytic ($k_y$) and conversion rate ($k_{ii1}$) in case of scheme-V.**

For case $T_1$, the simulated time profiles of the product ($X$) and product ($Y$), when [C-D] events (4 [C-D]) fine-tune the intermediate rate $k_{ii1}$ and the catalytic rate $k_y$ either individually (solid lines in **A, B, C, and D**). Time evolution of **E)** product ($X$) and **F)** product ($Y$), when [C-D] event (4 [C-D]) alters the intermediate rate $k_{ii1}$ and the catalytic rate $k_y$ collectively (solid lines). The simulation has been performed by taking the binding rates similar and keeping all other rate constants the same as in Table A5.5. The dashed lines represent the model-predicted profile for CT, and the dotted lines indicate the model-predicted profile altering the length ($k_{b1}/L$, $k_{b2}/L$, $L = 0.1$) at different [C-D] cycles.

#### Text S9 Observation of the kinetics of the biochemical reaction following mechanisms like scheme-VI

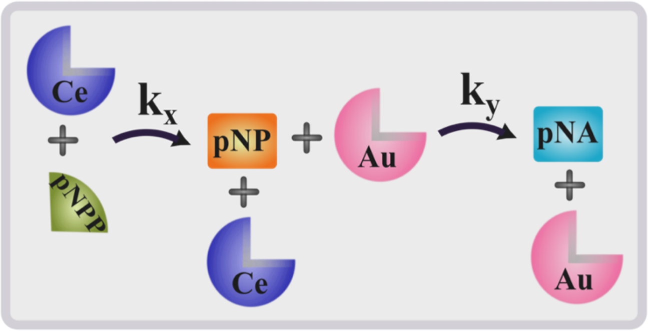

**Fig. A5.20 Probable schematic of heterogeneous catalysis by ceria (Ce) and gold (Au) particle-based enzyme mimicking materials.**

**Table S14 Abbreviation of different variables involved in the scheme-VI**

| Number | Symbol | Description ( in Conc., $\mu M$) |
|---|---|---|
| 1 | $E_1$ | The free amount of ceria nano-particle (first enzyme, Ce) |
| 2 | $S$ | The free amount of p-nitro phenyl phosphate (substrate, pNPP) |
| 3 | $E^1S$ | The enzyme-substrate complex formed from the binding of p-nitro phenyl phosphate (substrate, $S$) with ceria nano-particle (enzyme, $E_1$) |
| 4 | $E^{\#1}S$ | Another intermediate state is produced from $E^1S$ |
| 5 | $X$ | The amount of p-nitro phenolate ion (product, pNP) |
| 6 | $E_2$ | The free amount of gold nano-particle (second enzyme, Au) |
| 7 | $E^2X$ | The enzyme-substrate complex formed from binding of p-nitro phenolate ion (product, $X$) with gold nano-particle (enzyme, $E_2$) catalyzed by $NaBH_4$ |
| 8 | $E^{\#2}X$ | Another intermediate state is produced from $E^2X$ |
| 9 | $Y$ | The amount of p-nitro phenolate amine (product, |

| | | pNA) |
|---|---|---|
| **Number** | **Reaction** | **The propensity of the reaction in $j^{th}$ compartment** |
| **1** | $E_1(j) + S(j) \rightarrow E^1S(j)$ | $a\_nu(1)_j = k_{b1} \times S(j) \times E_1(j)$ |
| **2** | $E^1S(j) \rightarrow E_1(j) + S(j)$ | $a\_nu(2)_j = k_{d1} \times E^1S(j)$ |
| **3** | $E^1S(j) \rightarrow E^{\#1}S(j)$ | $a\_nu(3)_j = k_{i1} \times E^1S(j)$ |
| **4** | $E^{\#1}S(j) \rightarrow E^1S(j)$ | $a\_nu(4)_j = k_{i2} \times E^{\#1}S(j)$ |
| **5** | $E^{\#1}S(j) \rightarrow X(j) + E_1(j)$ | $a\_nu(5)_j = k_x \times E^{\#1}S(j)$ |
| **6** | $E_2(j) + X(j) \rightarrow E^2X(j)$ | $a\_nu(6)_j = k_{b2} \times E_2(j) \times X(j)$ |
| **7** | $E^2X(j) \rightarrow E_2(j) + X(j)$ | $a\_nu(7)_j = k_{d2} \times E^2X(j)$ |
| **8** | $E^2X(j) \rightarrow E^{\#2}X(j)$ | $a\_nu(8)_j = k_{ii1} \times E^2X(j)$ |
| **9** | $E^{\#2}X(j) \rightarrow E^2X(j)$ | $a\_nu(9)_j = k_{ii2} \times E^{\#2}X(j)$ |
| **10** | $E^{\#2}X(j) \rightarrow Y(j) + E_2(j)$ | $a\_nu(10)_j = k_y \times E^{\#2}X(j)$ |
| **11** | $S(j) \rightarrow S(j+1)$ | $a\_nu(11)_j = D_h \times S(j), j = 1\ to\ (K-1)$ |
| **12** | $S(j) \rightarrow S(j-1)$ | $a\_nu(12)_j = D_h \times S(j), j = 2\ to\ K$ |
| **13** | $X(j) \rightarrow X(j+1)$ | $a\_nu(13)_j = D_h \times X(j), j = 1\ to\ (K-1)$ |
| **14** | $X(j) \rightarrow X(j-1)$ | $a\_nu(14)_j = D_h \times X(j), j = 2\ to\ K$ |
| **15** | $Y(j) \rightarrow Y(j+1)$ | $a\_nu(15)_j = D_h \times Y(j), j = 1\ to\ (K-1)$ |
| **16** | $Y(j) \rightarrow Y(j-1)$ | $a\_nu(16)_j = D_h \times Y(j), j = 2\ to\ K$ |
| **Algebraic Relation** | | |
| $E1_t(j) = E_1(j) + E^1S(j) + E^{\#1}S(j)$ | | |
| $E2_t(j) = E_2(j) + E^2X(j) + E^{\#2}X(j)$ | | |
| $S_t(j) = S(j) + E^1S(j) + E^{\#1}S(j) + X(j) + E^2X(j) + E^{\#2}X(j) + Y(j)$ | | |

**Table S15 Description of the parameters associated with the Scheme-VI for three different cases**

| Symbol | Description | Unit | [T1] | [T2] | [T3] |
|---|---|---|---|---|---|
| $k_{b1}$ | Binding rate of the Ceria nano-particle ($E_1$) with p-nitro phenyl phosphate ($S$) | $\mu M^{-1}min^{-1}$ | 0.002 | 0.002 | 0.002 |
| $k_{d1}$ | The dissociation rate of the | $min^{-1}$ | 10 | 10 | 10 |

| | | | | | |
|---|---|---|---|---|---|
| | enzyme-substrate complex $E^1S$ | | | | |
| $k_{i1}$ | The conversion rate of the enzyme-substrate complex $(E^1S)$ to another intermediate conformer $(E^{\#1}S)$ | $\mu M^{-1}min^{-1}$ | 2.0 | 2.0 | 0.002 |
| $k_{i2}$ | The conversion rate of intermediate conformer $E^{\#1}S$ to enzyme-substrate complex $(E^1S)$ | $min^{-1}$ | 0.1 | 0.1 | 10 |
| $k_x$ | Catalytic rate or formation rate of p-nitro phenolate ion $(X)$ from $(E^{\#1}S)$ | $min^{-1}$ | 1.0 | 1.0 | 4.0 |
| $k_{b2}$ | Binding rate of the Gold nano-particle $(E_2)$ with p-nitro phenolate ion $(X)$ | $\mu M^{-1}min^{-1}$ | 0.006 | 0.006 | 0.002 |
| $k_{d2}$ | The dissociation rate of the enzyme-substrate complex $(E^2X)$ | $min^{-1}$ | 10.0 | 10.0 | 10 |
| $k_{ii1}$ | The conversion rate of the enzyme-substrate complex $(E^2X)$ to another intermediate conformer $(E^{\#2}X)$ | $\mu M^{-1}min^{-1}$ | 2.0 | 2.0 | 0.002 |
| $k_{ii2}$ | The conversion rate of the intermediate conformer $(E^{\#2}X)$ to enzyme-substrate complex $(E^2X)$ | $min^{-1}$ | 0.1 | 0.1 | 10 |
| $k_y$ | Catalytic rate or formation rate of p-nitro phenylamine$(Y)$ from $(E^{\#2}X)$ | $min^{-1}$ | 2.0 | 2.0 | 4.0 |
| $D_h$ | Forward/ Backward diffusion rate of the p-nitro phenyl phosphate $(S)$/ p-nitro phenolate ion $(X)$/ p-nitro phenylamine$(Y)$ | $cm^2min^{-1}$ | 0.023 | 0.023 | 0.023 |
| $S_t$ | The total amount of p-nitro phenyl phosphate $(S)$ | $\mu M$ | | | 1000 |
| $E_{1t}$ | The total amount of Ceria nano-particle $(E_1)$ | $\mu M$ | | | 2000 |
| $E_{2t}$ | The total amount of Gold nano-particle $(E_2)$ | $\mu M$ | | | 2000 |
| $K$ | Total number of compartments | - | | | 50 |

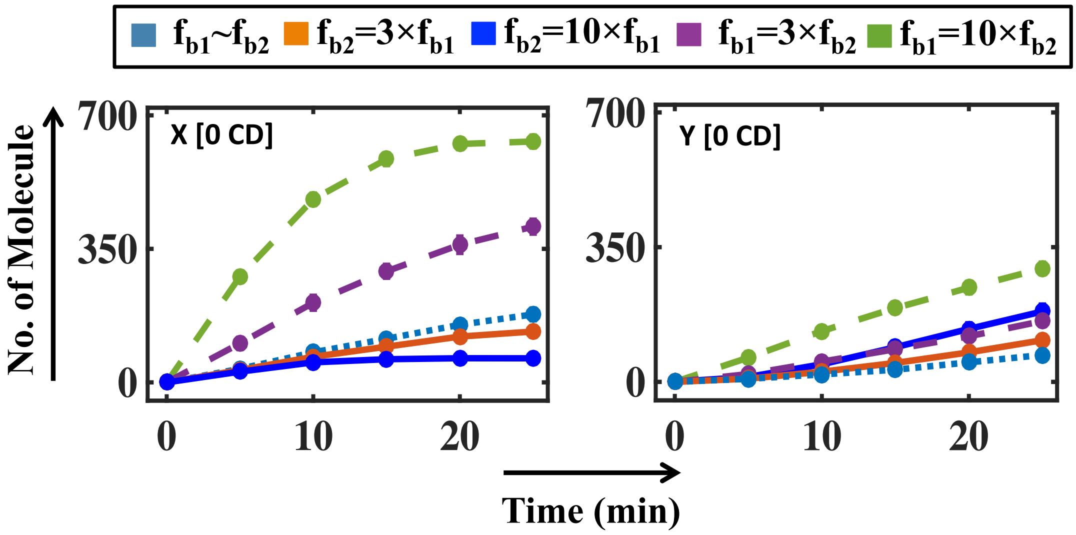

**Fig. S21 Simulated trajectory of the product altering the binding rate of the two enzymes($k_{b1}, k_{b2}$) in case of schematic-VI.**

The simulated trajectory of product X and product Y under varying the binding rate of two enzymes ($E1$, $E2$) in different combinations[$(i)$ $k_{b1}(0.002) \sim k_{b2}(0.002)$; $(ii)$ $k_{b1}(0.002) = 3 \times k_{b2}(0.006)$; $(iii)$ $k_{b1}(0.002) = 10 \times k_{b2}(0.02)$; $(iv)$ $k_{b2}(0.002) = 3 \times k_{b1}(0.006)$; $(v)$ $k_{b2}(0.002) = 10 \times k_{b1}(0.02)$; ] keeping all other rate constants the same as depicted in Table A5.7.

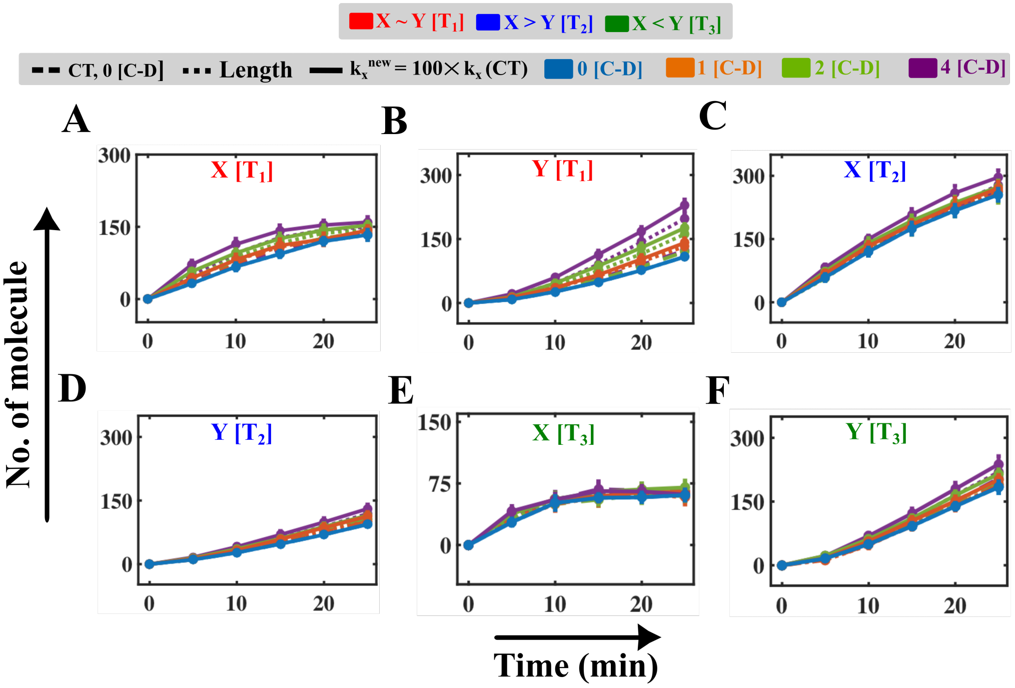

**Fig. S22 Simulated trajectory of the product altering the catalytic rate of enzyme E₁ $(k_x)$ in case of schematic-VI under different conditions.**

The simulated trajectory of product $X$ (**A**) case $T_1$, **C**) case $T_2$ and **E**) case $T_3$ ) and product $Y$ (**B**) case $T_1$, **D**) case $T_2$ and **F**) case $T_3$ ) by enhancing the catalytic rate of enzyme E₁ 100 times than WT case $(k_x(new) = 100 \times k_x(WT))$. The rest of the rate constant values are similar as depicted in **Table A5.7**.

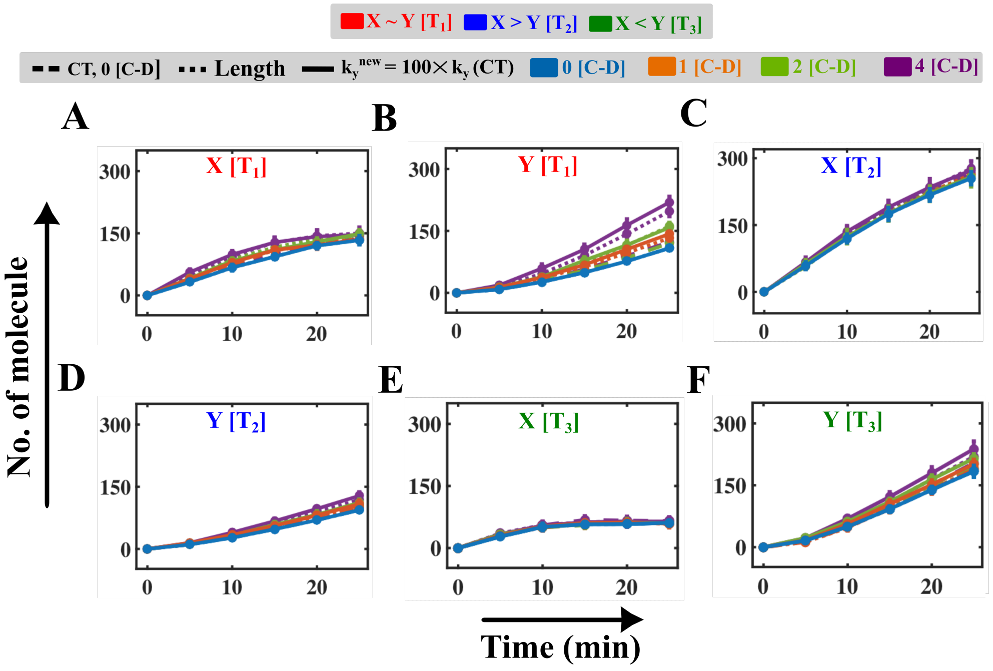

**Fig. S23 Simulated trajectory of the product altering the catalytic rate $(k_y)$ in case of schematic-VI under different conditions.**

The simulated trajectory of product $X$ (**A**) case $T_1$, **C**) case $T_2$ and **E**) case $T_3$ ) and product $Y$ (**B**) case $T_1$, **D**) case $T_2$ and **F**) case $T_3$ ) by enhancing the catalytic rate of enzyme E₁ 100 times than WT case $(k_y(new) = 100 \times k_y(WT))$. The rest of the rate constant values are similar as depicted in **Table A5.7**.

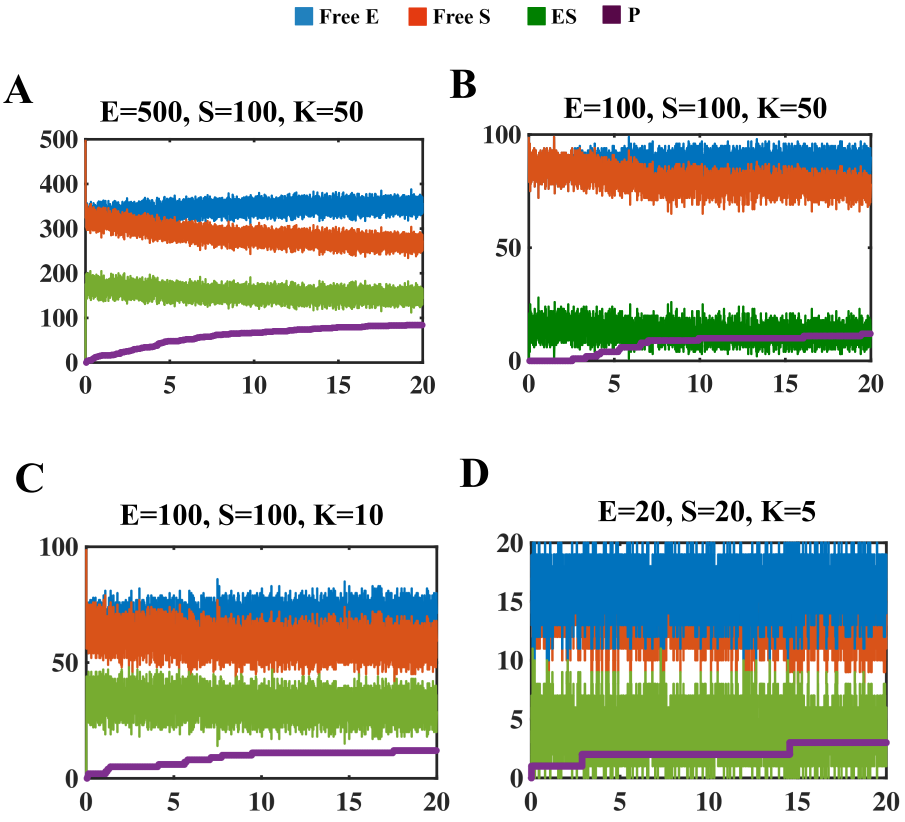

**Fig. S24 Effect of molecular fluctuations for very low number of enzyme and substrate**
Effect of molecular fluctuations on the dynamics of enzyme (E), substrate (S), enzyme-substrate complex (ES) and product (P) following a simple reaction. Time profiles of E, S, ES and P performing SRDA with NMCD for 1 [C-D] cycles. With decreasing the number of molecules, the noise increases. K denotes the number of compartments.
